# The Tgf-β family member Gdf6Y determines the male sex in *Nothobranchius furzeri* by suppressing oogenesis-inducing genes

**DOI:** 10.1101/2023.05.26.542338

**Authors:** Annekatrin Richter, Hanna Mörl, Maria Thielemann, Markus Kleemann, Raphael Geißen, Robert Schwarz, Carolin Albertz, Philipp Koch, Andreas Petzold, Marco Groth, Nils Hartmann, Amaury Herpin, Christoph Englert

## Abstract

The short-lived African killifish *Nothobranchius furzeri* lives in seasonal freshwater ponds and has evolved remarkable traits to survive in this limited environment. One of those traits is a genetic XX/XY sex-determination system, which ensures an equal distribution of both sexes. Comparisons of female and male genomic sequences identified the Y-chromosomal copy of the TGF-β family member *gdf6* as the candidate male sex-determining (SD) gene, which was named *gdf6Y* in contrast to the X-chromosomal allele *gdf6X*. CRISPR/Cas9-mediated inactivation of *gdf6Y* in *N. furzeri* led to a complete male-to-female sex reversal in XY animals. The homozygous inactivation of *gdf6X* on the other hand led to a detrimental phenotype post-hatching. This phenotype was compensated by *gdf6Y*, revealing that the latter became the SD gene while retaining at least some of its original *gdf6* function. *Gdf6Y* is expressed in testicular somatic cells already prior to hatching, where it represses the germ cell-intrinsic feminizing gene *foxl2l*. We have identified components of the TGF-β signaling pathway, especially the inhibitor of DNA binding genes *id1/2/3*, and the mRNA decay activator *zfp36l2*, as Gdf6Y targets. We conclude that Gdf6Y exerts its function as the male sex-determining gene by suppressing female-specific genes in the developing gonad of male *N. furzeri*.

## INTRODUCTION

The African turquoise killifish *Nothobranchius furzeri* is the captive vertebrate with the shortest lifespan of about 6 months reported so far (Platzer and Englert 2016). Despite its short lifespan, *N. furzeri* shows remarkable aging phenotypes and, therefore, is an upcoming model organism, especially in aging research (Kim et al. 2016; Reuter et al. 2018). The annual killifish species inhabits ephemeral ponds in the South of Zimbabwe and Mozambique. From several of these isolated wild populations, laboratory strains with varying lifespans and different tail colors were derived e. g., the yellow-tailed strain GRZ with a lifespan of 6 months from Zimbabwe (Jubb 1971) and the strains MZM0403 and MZM0410 with an around 1-year lifespan from Mozambique (Terzibasi et al. 2008; Tozzini et al. 2013). In the wild, the embryos survive the dry season by facultative developmental arrests named diapauses. In direct development without diapause, the embryos enter the pharyngula stage and continue organogenesis at about 9 days post-fertilization (dpf), which is completed at around 14 dpf (Api et al. 2018). Between three and four weeks after fertilization, the embryos are ready to hatch. This so-called hatching stage (0 days post-hatching – 0 dph) can be maintained for several weeks. After hatching the larvae quickly grow, reach sexual maturation, and develop a strong sexual dimorphism at three to four weeks post-hatching. With a 1:1 sex ratio, males were found to be the heterogametic sex, and an XY sex-determination system was suggested (Valenzano et al. 2009). Only when the species’ genome was sequenced, the sex chromosome pair was identified (Reichwald et al. 2015; Valenzano et al. 2015) and the Y-chromosomal copy of the growth differentiation factor 6 gene (*gdf6*), which was therefore named *gdf6Y*, was proposed as the potential sex-determining (SD) gene (Reichwald et al. 2015).

Gdf6 is a secreted ligand-protein that belongs to the bone morphogenetic proteins (BMPs), a group of the transforming growth factor beta (TGF-β) superfamily. These proteins consist of an N-terminal prodomain, which is proteolytically cleaved-off upon maturation, and the C-terminal ligand portion, which binds as a dimer to members of the corresponding TGF-β receptor family. This leads to the intracellular activation of receptor-regulated SMAD transcription factors (R-SMADs) by phosphorylation and ultimately regulates gene expression. Other TGF-β ligand and receptor family members, including orthologs of *amh*, *amhrII*, *bmprI*, and *gsdf*, have been previously identified as SD genes in various fish species (Hattori et al. 2012; Kamiya et al. 2012; Myosho et al. 2012; Kikuchi and Hamaguchi 2013; Li et al. 2015; Pan et al. 2019; Feron et al. 2020; Peichel et al. 2020; Herpin et al. 2021; Pan et al. 2021; Holborn et al. 2022). Therefore, factors involved in TGF-β signaling constitute one functional group, out of which SD genes are recruited in fishes. The other main group of SD gene candidates is transcription factors, especially those already involved in sex determination like *Dmrt1* and *Sox* genes (Pan et al. 2021). The SD gene of the rainbow trout (*Oncorhynchus mykiss*), *sdY*, is an exception, as its ancestral gene *irf9* is an immune-related transcription factor (Yano et al. 2012).

In the sequenced *N. furzeri* strain GRZ, *gdf6Y* differs from the X chromosomal *gdf6* copy, hereafter called *gdf6X*, in 22 single nucleotide polymorphisms (SNPs) and a 9-bp deletion in the coding sequence (CDS) of the prodomain (Fig. 1 A; Supplemental Fig. S1 A; Reichwald et al. (2015)). While the non-synonymous SNPs and the 9-bp deletion are conserved among the laboratory *N. furzeri* strains analyzed so far, the size of the genomic region, which accumulated differences between the X and Y chromosome, varies greatly from 196 kb to 37 Mb in the strains MZM0403 and MZM0410, respectively, indicating the ongoing evolution of the species’ sex chromosomes. In other teleost and tetrapod species, including zebrafish, *Xenopus laevis*, mice, and humans, *GDF6* orthologs were predominantly shown to be involved in joint development and eye formation (Settle et al. 2003; Hanel and Hensey 2006; Asai-Coakwell et al. 2007; Tassabehji et al. 2008; Asai-Coakwell et al. 2009; Gosse and Baier 2009; den Hollander et al. 2010; Reed and Mortlock 2010; Asai-Coakwell et al. 2013), but also the involvement in vascularization (Crosier et al. 2002; Hall et al. 2002; Krispin et al. 2018) and melanoma as well as melanocyte development were observed (Venkatesan et al. 2018; Gramann et al. 2019).

**Figure 1.**
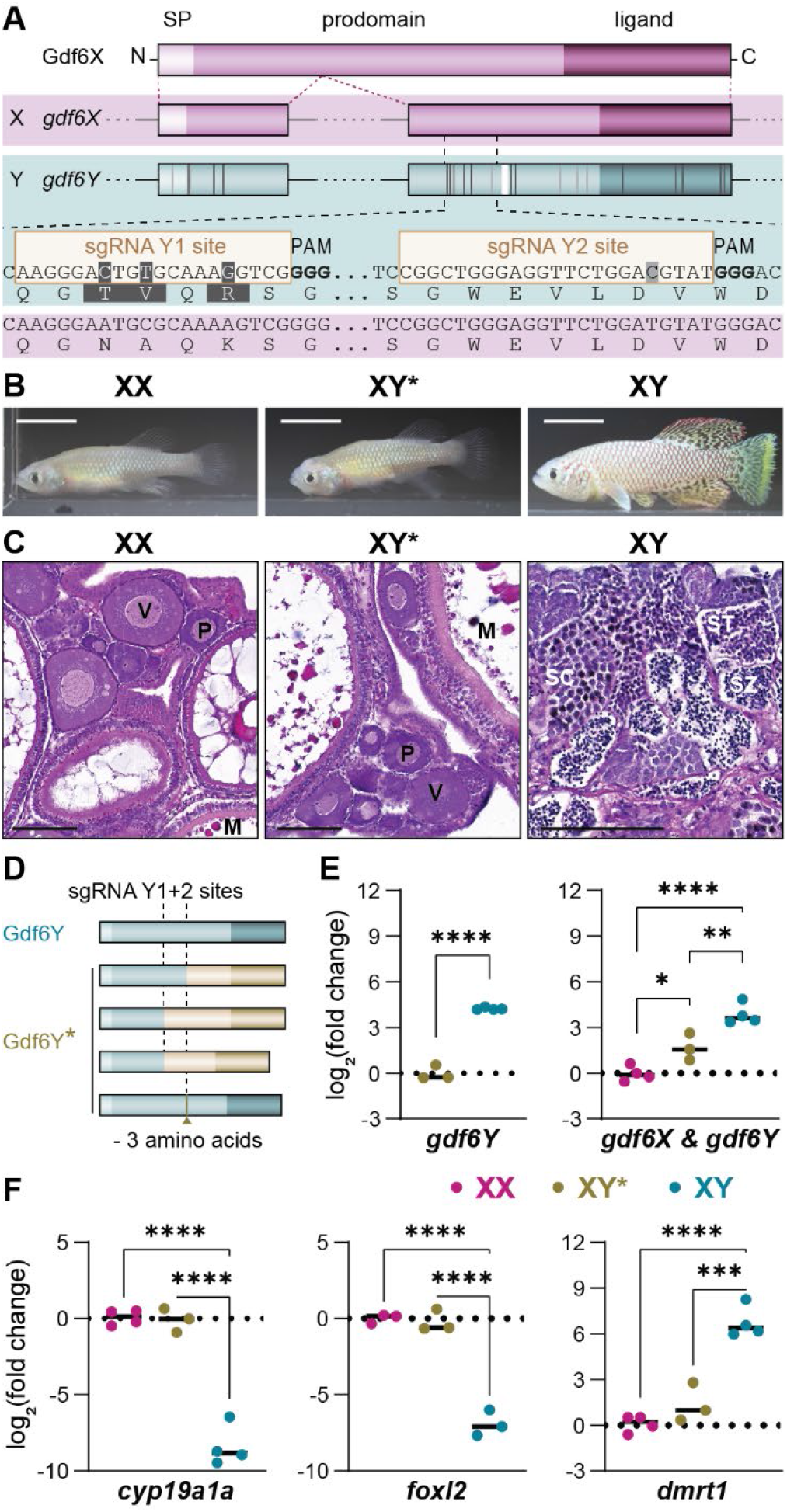
The inactivation of *gdf6Y* leads to full male-to-female sex reversal in *Nothobranchius furzeri*. *(A)* Schematic of the Gdf6X protein domains (SP – signal peptide), the exonal distribution and comparison of the coding sequences of *gdf6X* and *gdf6Y* as well as the mutation strategy with two *gdf6Y*-specific sgRNAs that facilitate DNA double-strand breaks at a distance of 109 bp. Vertical lines in the *gdf6Y* exons indicate coding sequence differences between *gdf6X* and *gdf6Y*. SNPs: Dark gray – non-synonymous, light gray – synonymous. White – 9 bp deletion. *(B)* The phenotype of mosaic *gdf6Y*^-^ animals (XY*, phenofemales) compared to an unaltered female (XX) and a wild-type male (XY) at about 3 months of age. Scale bar: 10 mm. *(C)* H&E staining of XX, XY*, and XY gonads at about 5 months of age. Oocytes: P – previtellogenic, V – vitellogenic, M – mature. SC – spermatocytes, ST – spermatids, SZ – spermatozoa. Scale bar: 100 µm. *(D)* Gdf6Y* variants that lead to full male-to-female sex reversal contain either a frameshift at one sgRNA position, a deletion between both sgRNAs, or a 9 bp deletion at the sgRNA Y2 site. *(E, F)* Expression of *gdf6X* and *gdf6Y* together and *gdf6Y* separately *(E)* as well as male and female marker genes *(F)* on mRNA-level in gonads of animals of the 1^st^ filial generation at about 2.4 months of age. XY* – phenofemales with frameshift mutations. Statistical testing by either a two-tailed, unpaired *t*-test (*gdf6Y*) or a 1-way ANOVA and Tukey’s post hoc test (others), (*) *P*<0.05, (**) *P*<0.01, (***) *P*<0.001, (****) *P*<0.0001.

Here, we demonstrate *gdf6Y* as the SD gene in *N. furzeri* by specifically disrupting the Y-chromosomal allele. The respective mutants showed a complete male-to-female sex reversal. The homozygous inactivation of the X-chromosomal allele *gdf6X* on the other hand led to a detrimental curled tail phenotype, which abolished proper larval swimming immediately after hatching. However, *gdf6Y* compensates for the loss of *gdf6X* and, therefore, must have additionally acquired the SD gene function. In this role, *gdf6Y* is expressed in the male gonad by a subset of somatic cells at 0 dph and in Sertoli cells in adulthood. At 0 dph, however, oogenesis has already begun in the female gonads, and thereby the sex in *N. furzeri* must be determined beforehand. As suggested by the analyses of mutant and transgenic mosaic animals, *gdf6Y* most likely determines the male sex by inhibiting the female gonadal development, for example by suppressing the expression of the oogenesis-initiating germ cell (GC) factor *foxl2l* (Nishimura et al. 2015; Kikuchi et al. 2019; Kikuchi et al. 2020). This is probably facilitated by genes directly regulated in response to Gdf6Y e.g., *id1* and *zfp36l2*, which we identified in a cell culture-based approach.

## RESULTS

### The disruption of *gdf6Y* leads to full male-to-female sex reversal

Being the only annotated gene at the peak of Y-specific sequence variation in the *N. furzeri* strain with the smallest SD region (SDR; MZM0403) and the gene with the strongest signal of positive selection in the SDR of the hereafter used, genome-sequenced strain GRZ, *gdf6Y* was proposed as the SD gene in *N. furzeri* (Reichwald et al. 2015). To test whether *gdf6Y* is indeed the male sex determinant in *N. furzeri*, we employed Cas9 and two sgRNAs specifically targeting three non-synonymous or one synonymous SNP in the 2^nd^ exon of *gdf6Y* (Fig. 1 A). This approach led to the highly unlikely result of 16 female and no male fish (*P* = 1.53 × 10^−5^; Fig. 1 B). Eight out of these sixteen females had XY chromosomes with an average mutation rate of 97 % in *gdf6Y* only, as determined by cloning the heterogeneous PCR products derived from the target locus and sequencing several clones (Supplemental Fig. S1 B, C, D). In contrast, the eight animals with XX chromosomes had no mutations in *gdf6X* that were detectable with either a T7 endonuclease I assay or by sequencing the PCR product (Supplemental Fig. S1 B, E, F), indicating that the inactivation was very efficient and highly specific. The thus generated, phenotypically female animals with XY chromosomes and an impaired *gdf6Y* are hereafter called phenofemales (XY*). The mosaic phenofemales had gonads, which were histologically indistinguishable from female ovaries (Fig. 1 C) and were fertile. Interestingly, either different frameshift mutations or an in-frame 9 bp deletion in the CDS of *gdf6Y*’s prodomain consistently resulted in the same phenotype (Fig. 1 D; Supplemental Tab. S1). Furthermore, the phenofemales had significantly reduced *gdf6Y* transcript levels compared to males and showed no difference from females in the gonadal expression of sex-specific marker genes (Fig. 1 E, F). Taken together, the inactivation of *gdf6Y* led to full male-to-female sex reversal, which confirms *gdf6Y* as the major male sex determinant in *N. furzeri*.

### YY* embryos show anomalies in eye and body axis development

According to Mendelian ratios, we expected animals with two Y-chromosomes (YY*) among the offspring of phenofemales and males (Fig. 2 A). As their adult progeny consisted only of males (XY), females (XX), and phenofemales (XY*) in equal ratio, we monitored the development of the phenofemales’ clutches and observed malformations of the eye and tail within the egg during organogenesis in about one-quarter of the embryos (Fig. 2 A; Supplemental Fig. S2 A). Those embryos died before hatching and were identified as the YY* animals (Supplemental Fig. S2 B). The YY* typical malformations were characterized by a lacking tail fin development and pigmentation as well as an incomplete closure of the optic fissure leading to a coloboma-like phenotype (Fig. 2 A; Supplemental Fig. S2 C). A histological comparison with normally developing animals at 0 dph also revealed a decreased thickness of retinal layers in the YY* embryos’ eyes (Supplemental Fig. S2 D).

**Figure 2.**
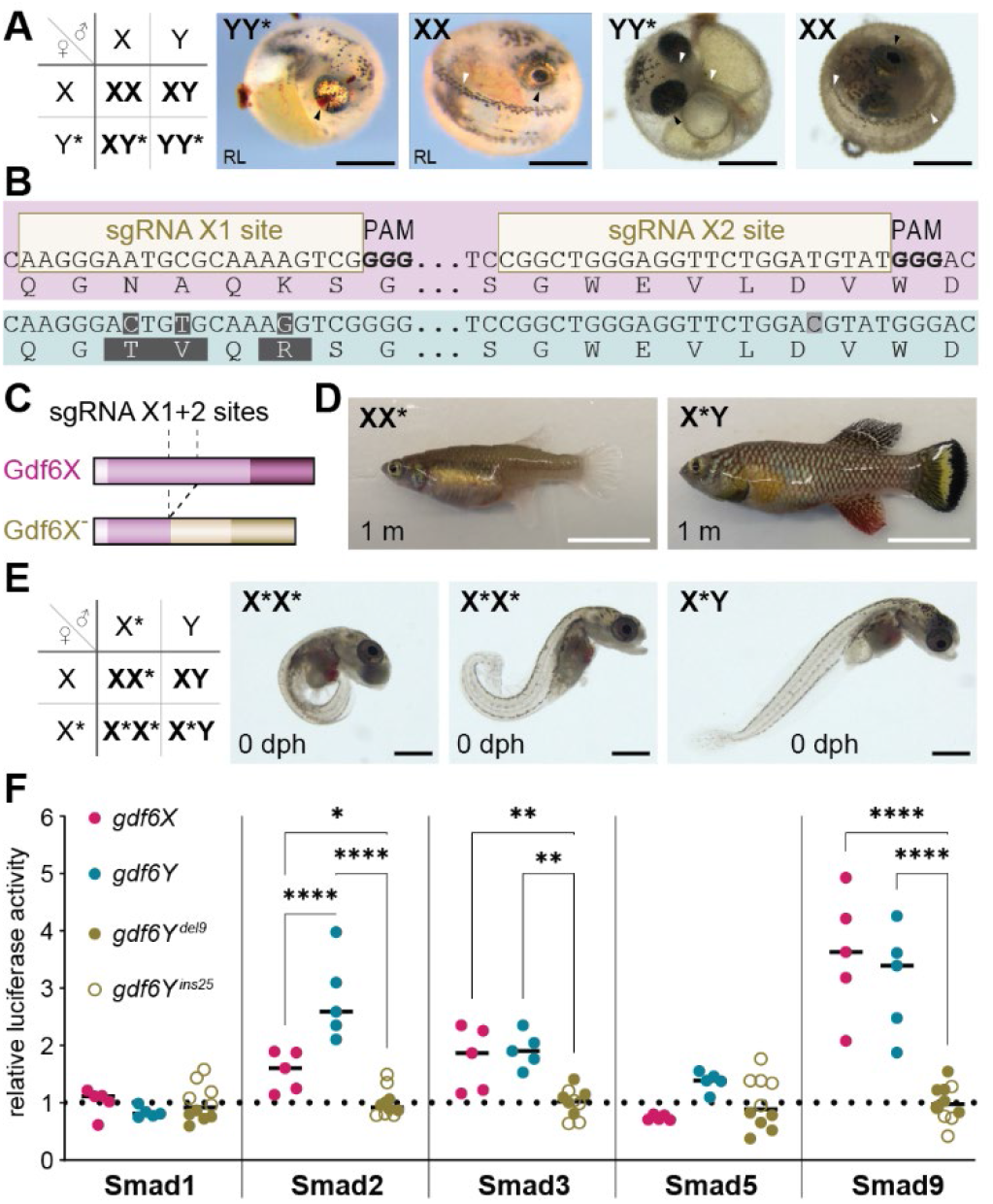
Gdf6Y can functionally cover for Gdf6X. *(A)* One quarter of the offspring from phenofemales (XY*) and males (XY) carries two Y-chromosomes (YY*), does not survive until hatching, and shows severe malformations within the egg compared to normally developed animals (here: XX at 0 dph). Shown are eggs from two different *gdf6Y*-mutants (left – *gdf6Y^del6, del8^*, right – *gdf6Y^del9^*). Arrows: Black – eye, white – tail. RL – reflected light microscopy. *(B)* Schematic of the *gdf6X* inactivation and *(C)* the obtained open reading frame disruption. *(D)* The phenotype of heterozygous *gdf6X*^-^ females (XX*) and males (X*Y). 1 m – 1 month. *(E)* Homozygous *gdf6X*^-^ (*gdf6X^del113^*) hatchlings (X*X*; n = 15) have a detrimental curled phenotype (two manifestations shown), which prevents regular swimming behavior, while siblings (X*X, X*Y, XY; n = 34) are normal. X*Y shown for comparison. *(A, E)* Pictures of pre-or post-hatchlings acquired by brightfield microscopy if not indicated otherwise. *(A, C, E)* Scale bars: Black – 500 µm, white – 10 mm. *(F)* Luciferase reporter assay to test the differential activation of Smad1, 2, 3, 5, and 9 upon expression of *gdf6X* or *gdf6Y* compared to *gdf6Y* mutant variants (*gdf6Y^del9^* and *gdf6Y^ins25^*). *Gdf6Y^del9^* has an in-frame 9 bp deletion in the prodomain’s CDS, while *gdf6Y^ins25^* harbors a 25 bp insertion at the same position leading to a frameshift and early Stop-codon. Statistical testing by a 2-way ANOVA and Holm-Šídák’s multiple comparisons tests, (*) *P*<0.05, (**) *P*<0.01, (****) *P*<0.0001.

As *GDF6* orthologs are known to be involved in eye, joint, and melanocyte formation in various species (Settle et al. 2003; Asai-Coakwell et al. 2009; den Hollander et al. 2010; Gramann et al. 2019), we wondered whether the lack of *gdf6X* in YY* embryos contributes to their malformations. To address this question, we employed Cas9 and two sgRNAs targeting the same positions previously used to disrupt *gdf6Y* but on the X-chromosome (Fig. 2 B). Using the Synthego ICE (Inference of CRISPR Editing) analysis tool (v3), we estimated an average mutation rate of 97 % and 14 % in *gdf6X* and *gdf6Y*, respectively, indicating that the inactivation of *gdf6X* was as efficient but not as specific as in the case of *gdf6Y* (Supplemental Fig. S2 E). Hence, we observed phenotypes in about one-third of the mosaic males and two-thirds of the mosaic females (Supplemental Fig. S2 F). Two out of ten mosaic males had one-sided microphthalmia, a probably clonal phenotype that was not passed on to subsequent generations. Furthermore, six out of nine mosaic females and one male did not straighten their body axes after hatching, which lead to a curled tail that disabled normal swimming. This detrimental phenotype was also found at varying degrees in all analyzed homozygous hatchlings (X*X*) of the isolated *gdf6X* frameshift mutant line (Fig. 2 C, E; Supplemental Tab. S1) and *gdf6X*/*gdf6Y* double mutant offspring (X*Y*) of phenofemales and mosaic *gdf6X* mutant males (Supplemental Fig. S2 G). However, this curled phenotype was fully compensated not only in female (XX*) but also in male (X*Y) heterozygous animals (Fig. 2 D), indicating that *gdf6Y*, aside from its SD function, can functionally compensate *gdf6X* inactivation. Thus, the lack of *gdf6X* does not contribute to the phenotype of the YY* embryos, which have one functional *gdf6Y* copy.

As the phenofemales were generated in the genome-sequenced strain GRZ, the detrimental phenotype of the YY* embryos could be caused by defects in or a dysregulation of the 339 genes annotated in the strain’s 26.1 Mb large SDR (sgr05: 15,031,832–41,162,746; Reichwald et al. (2015)). To investigate whether genes in the SDR have a sex chromosome-specific and copy number-dependent regulation, previously published transcriptome data from male and female whole embryos at 10 dpf were consulted (Reichwald et al. 2015). The expression of the most significantly up- and downregulated genes between males and females within the SDR was analyzed in phenofemales and YY* embryos at 11 dpf (Supplemental Fig. S2 H). We found that the most significantly upregulated gene in females, *rnf19a*, indeed showed an X chromosome-dependent expression, while the most downregulated gene, *pctp*, showed an increased expression only in the presence of a Y chromosome. Thus, the sequence changes accumulating on the Y chromosome can lead to both a locally enhanced and reduced gene expression, potentially interfering with the development of YY* embryos.

Given that *gdf6X* and *gdf6Y*, apart from SD, seem to be functionally equivalent, we assessed the signaling activities of the respective proteins using a heterologous cellular reporter system (Herpin et al. 2021). To this end, medaka cells harboring different Gal4-Smad fusion proteins (Smad1, 2, 3, 5, and 9) and a responsive luciferase reporter were transfected with expression plasmids for *gdf6X* and *gdf6Y* as well as two *gdf6Y* mutants as controls. Compared to the two *gdf6Y* mutant variants both *gdf6X* and *gdf6Y* showed similar signaling properties by significantly inducing the activation of Smad2, 3, and 9 (Fig. 2 F). Notably, Smad2 activation by Gdf6Y led to significantly higher luciferase activity than activation by Gdf6X. Therefore, Gdf6Y’s SD function did probably not evolve by an entire switch of the involved TGF-β signaling partners but could have been promoted by enhanced Gdf6Y signaling capabilities.

### A subset of the testicular somatic supporting cells expresses *gdf6Y*

As *gdf6Y* covers the functions of *gdf6X* and additionally determines the male sex in *N. furzeri*, we were interested in how it governs sexual development. It was previously shown that *gdf6Y* is expressed in adult testes while *gdf6X* is not expressed in adult gonads (Reichwald et al. (2015); Supplemental Fig. S3 A). To address the cellular localization of *gdf6Y*’s testicular expression, we used RNAscope as a very sensitive and specific *in situ* hybridization technique. Applying a *gdf6Y*-specific probe together with probes for the GC marker *ddx4* and one of the somatic cell markers *amh*, *dmrt1*, or *wt1a* (Kobayashi et al. 2004; Kluver et al. 2007; Kluver et al. 2009), we found that *gdf6Y* is co-expressed with all three somatic cell markers in proximity to the spermatogonia of the adult testes (Fig. 3 A; Supplemental Fig. S3 B, C). Next, we analyzed the gonadal expression of *gdf6Y* at 0 dph. Marking the GCs with *ddx4* and the somatic cells with *amh*, *dmrt1*, or *wt1a*, we observed that *gdf6Y* is expressed in a subset of the latter in the still largely undifferentiated male gonads (Fig. 3 B; Supplemental Fig. S3 D, E). In contrast to this, the female gonads at 0 dph, where *gdf6Y* is not expressed, are larger and more differentiated due to a greater cell number and already developing oocytes, respectively (Supplemental Fig. S3 F). Taken together, *gdf6Y* expression in both the developing as well as the adult testes was observed in somatic supporting cells.

**Figure 3.**
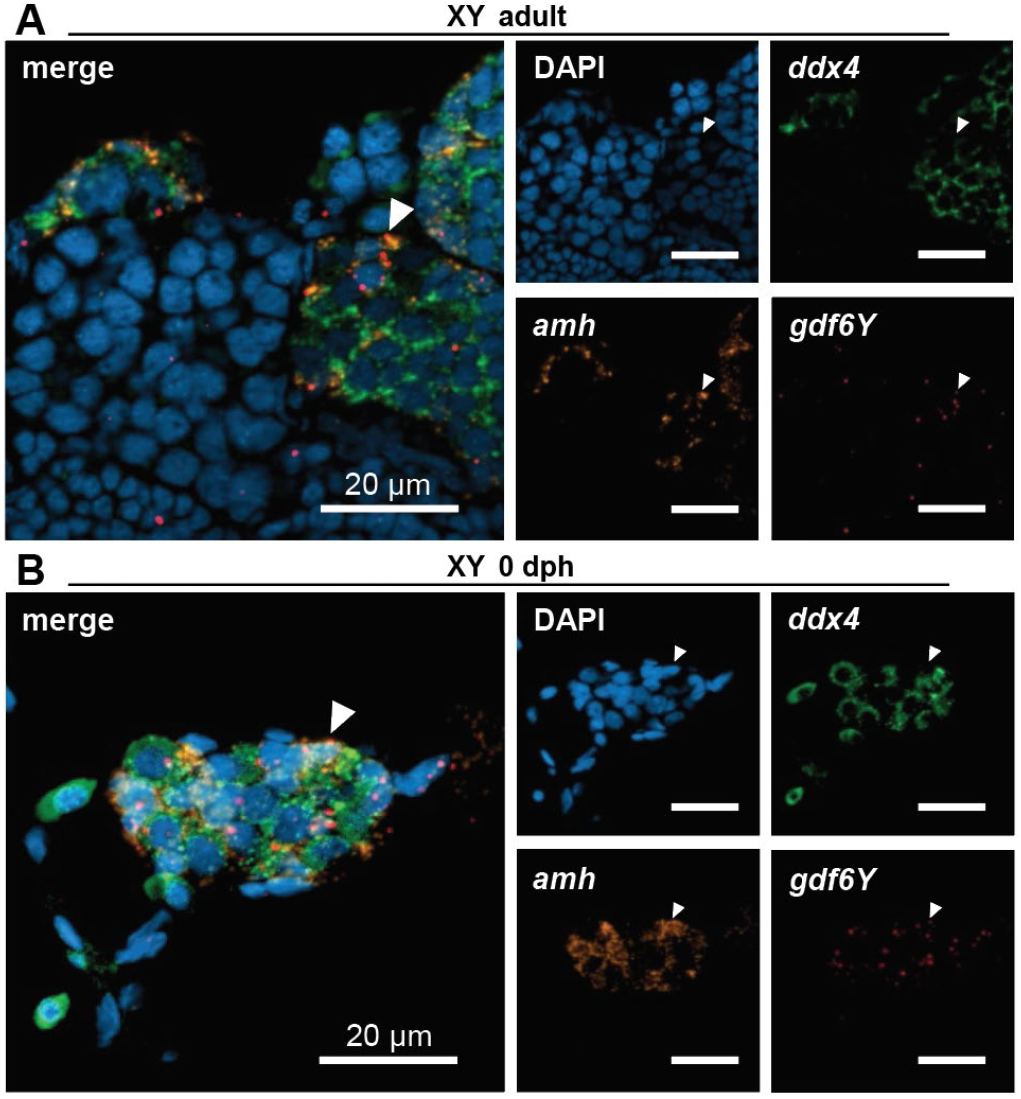
Localization of the gonadal *gdf6Y* expression in XY animals. Transcripts of *gdf6Y* (red) and the GC marker *ddx4* (green) as well as the *(A)* Sertoli or *(B)* somatic supporting cell marker *amh* (orange) were detected at *(A)* adult and *(B)* 0 dph stage. *(A, B)* White arrows – cells co-expressing *gdf6Y* and *amh*.

In contrast to *gdf6Y* and *gdf6X*, their upstream neighbor *sybu* is expressed in oocytes of the developing and adult ovaries (Supplemental Fig. S3 G, H), where its mRNA localizes to the vegetal pole contributing to dorsal determination of the embryo (Nojima et al. 2010; Colozza and De Robertis 2014; Oh and Houston 2017). To analyze DNA methylation as a potential cause of the differential expression of *gdf6X* and *gdf6Y* as well as *sybu* in adult gonads, bisulfite sequencing of the region between *gdf6X/Y* and *sybu* was performed on gonadal tissues from males, females, and phenofemales (Supplemental Fig. S4 A, B). While the identified DNA methylation patterns were highly symmetrical on both DNA strands, we found that neither the promoter of *gdf6X* nor *gdf6Y* was methylated in any of the analyzed sexes (Supplemental Fig. S4 A, B). Conversely, the *sybu* promoter was hypomethylated on the X but hypermethylated on the Y chromosome, which translated to a strictly X-chromosomal expression in testes (Supplemental Fig. S4 B, C). In phenofemales however, the methylation of the *sybu* promoter on the Y chromosome decreased compared to males (Supplemental Fig. S4 B). Therefore, *sybu* transcripts from the Y chromosome were also sequence verified in phenofemale ovaries (Supplemental Fig. S4 C). An explanation for the altered Y-chromosomal methylation in phenofemales could be demethylation in the maternal germline and the adoption of the methylation pattern from the paternal X chromosome, as passive maternal demethylation and a take-over of the paternal methylome was shown in zebrafish (Jiang et al. (2013); reviewed in Ortega-Recalde and Hore (2019)). This implies that the silenced genes of the Y chromosome are reactivated in phenofemales. The intergenic region between *sybu* and *gdf6X* or *gdf6Y*, which contains two Y-specific transposable elements (TEs; Supplemental Fig. S4 A) was strongly methylated at all analyzed positions, suggesting a transcriptional insulation of the two promoters. Notably, we found that *sybu* transcript variants are sex-specific in *N. furzeri* gonads with the shortest one (NCBI Reference Sequence: XM_015949384.2), which has an alternative transcription start site, being ovary-specific (Supplemental Fig. S4 C). In conclusion, upstream methylation does not seem to control the differential testicular expression of *gdf6X* and *gdf6Y*.

### Gdf6Y suppresses the onset of female development prior to hatching

To understand the ongoing molecular processes in the developing gonads, we analyzed previously published transcriptome data (Reichwald et al. 2015). Differentially expressed genes (DEGs) between males and females were identified at three different developmental stages, namely the pharyngula stage during organogenesis at 10 dpf, the hatching stage (0 dph), and the larval stage at 3 dph. In the whole embryo at 10 dpf, 3 transcripts were already noticeably upregulated in males and continued to do so in all stages (Supplemental Fig. S5 A). In phenofemales, those transcripts were expressed like in males and, therefore, are not involved in *gdf6Y-*driven sex determination in *N. furzeri* (Supplemental Fig. S5 B). In male trunk parts, 147 and 133 genes were upregulated at 0 and 3 dph, respectively (Supplemental Fig. S5 A). In contrast, the number of upregulated genes in female trunks was higher and increased from 322 to 1079 from 0 to 3 dph, respectively (Supplemental Fig. S5 A). From the identified DEGs at 0 dph 35.4 % of the female-specific and 10.9 % of the male-specific transcripts were recalled at 3 dph (Supplemental Fig. S5 C). We performed an overrepresentation analysis (ORA) to obtain enriched KEGG (Kyoto Encyclopedia of Genes and Genomes) pathways for the upregulated genes in males and females at 0 and 3 dph. In males, the most enriched pathway was steroid biosynthesis at 0 dph (Supplemental Fig. S5 D). Expectedly, no significantly enriched pathways were identified at 3 dph given the little overlap of male-specific transcripts between the two time points (Supplemental Fig. S5 C). In females, pathways related to cell cycle, DNA repair, meiosis, and oocyte maturation were enriched at 0 and 3 dph despite the incomplete overlap of DEGs between both time points (Supplemental Fig. S5 C, E, F). This result validates our observation of an already occurring oogenesis in female gonads at 0 dph from the *in situ* hybridization experiments (Fig. 3 B; Supplemental Fig. S3 F). Therefore, the animals’ sex is already determined at 0 dph, while the maintenance of *gdf6Y* expression until adulthood suggests that it might be reversible.

In medaka, *foxl2l* is a GC-intrinsic factor that promotes oogenesis by inducing *rec8a* and suppressing spermatogenesis (Nishimura et al. 2015; Kikuchi et al. 2019; Kikuchi et al. 2020). At 0 and 3 dph, the *N. furzeri* ortholog of *foxl2l* and its putative target gene *rec8* were among the significant DEGs (Supplemental Fig. S5 A). In contrast to the female somatic cell marker *foxl2* and like in medaka, *foxl2l* is upregulated in females at 0 and 3 dph but not in adult gonads (Fig. 1 F; Fig. 4 A; Supplemental Fig. S6 A). Analyzing trunk parts at 0 dph and adult gonads, we showed that *foxl2l* is regulated the same way in phenofemales as it is in females, which places the gene downstream of *gdf6Y* (Fig. 4 B). Furthermore, RNAscope analyses confirmed a *foxl2l* expression in ovarian GCs at 0 dph by co-localization with *ddx4* (Fig. 4 C). We also observed *foxl2l* expression in testicular GCs in proximity to *gdf6Y* expressing somatic cells in one out of six analyzed male individuals at 0 dph (Fig. 4 D). This observation suggests that Gdf6Y could affect the expression in GCs, probably leading to *foxl2l* suppression. Searching for *foxl2l* expression foci in adult gonads, we found a single undifferentiated *foxl2l* positive GC in an ovary and none in testes (Supplemental Fig. S6 B). To verify that *foxl2l* influences SD in *N. furzeri* like in medaka, we employed Cas9 and two sgRNAs simultaneously targeting the single exon of *foxl2l* in males, females, and phenofemales. This approach led to various indels disrupting the gene with an efficiency of 63 % on average (Supplemental Fig. S6 C, D). Next, we analyzed the expression of sex-linked genes in the trunk parts of the mosaic *foxl2l* mutants at 0 dph. While neither *gdf6X* nor *gdf6Y* expression was changed in the mutants of all sexes, we observed a significant decrease of marker genes for cell division (*tacc3*) and oocyte development (*zp3/4*, *sybu*) to nearly male levels in the mutant females and phenofemales (Fig. 4 E; Supplemental Fig. S6 E, F). The aromatase gene *cyp19a1a* and the meiosis marker *rec8*, a putative target of *foxl2l*, decreased only in some but not all mutant females and phenofemales (Supplemental Fig. S6 F). Taking the mosaic nature of the mutants into account, this data supports that *foxl2l* promotes oogenesis in *N. furzeri* and acts downstream of *gdf6Y*.

**Figure 4.**
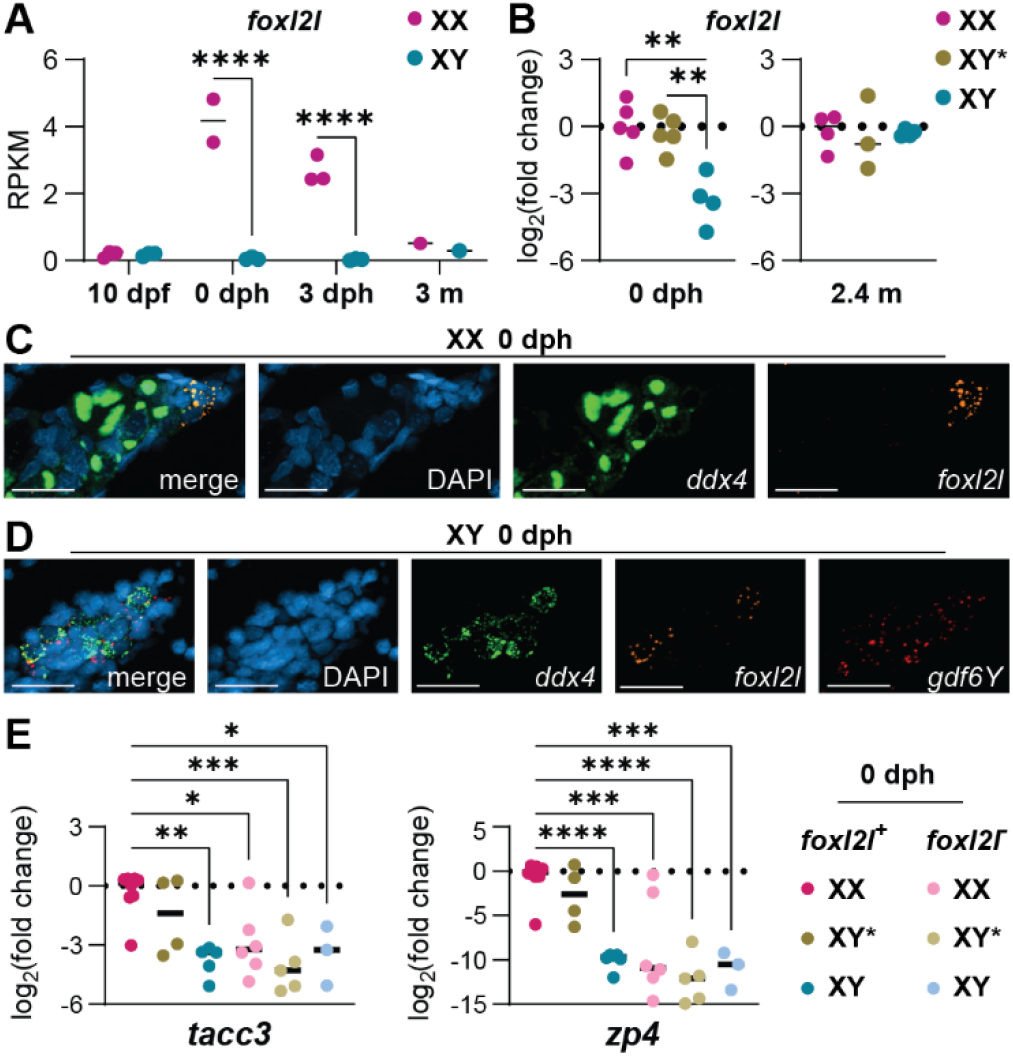
The *foxl2l* gene contributes to the female sex differentiation in *Nothobranchius furzeri* and is potentially repressed by *gdf6Y*. *(A*) Expression analysis of *foxl2l* by RNA-Seq in the whole embryo at 10 dpf, the trunk at 0 or 3 dph, and the gonad at 3 months (m) of age in XX and XY animals. RPKM – Reads per kb of transcript per Million mapped reads. 2-way ANOVA and Šidák’s post hoc test. *(B)* Expression of *foxl2l* on mRNA-level in trunks at 0 dph and adult gonads (2.4 m) of females (XX), phenofemales (XY*), and males (XY). *(C, D)* XX and XY gonadal expression at 0 dph of *foxl2l* (orange), the GC marker *ddx4* (green), and *(D) gdf6Y* (red). *(C, D)* Scale bar: 20 µm. *(E)* Expression of female marker genes on mRNA-level in trunks of females (XX), phenofemales (XY*), and males (XY) and their mosaic *foxl2l*^-^ counterparts at 0 dph. *(B, E)* 1-way ANOVA and Tukey’s post hoc test. *(A, B, E)* (*) *P*<0.05, (**) *P*<0.01, (***) *P*<0.001, (****) *P*<0.0001.

To investigate, if the presence of *gdf6Y* is sufficient to suppress oogenesis in *N. furzeri* development, we used Tol2 transgenesis to drive *gdf6Y* expression under the control of a ubiquitous *actb2* promoter from *D. rerio* (Supplemental Fig. S6 G; Hartmann and Englert (2012)). Analyzing the trunk parts of transgene-positive mosaic animals at 0 dph, we found that the expression of the female-specific genes *rec8*, *zp3/4*, and *sybu* was significantly decreased in transgenic XX animals compared to non-transgenic females (Supplemental Fig. S6 H). The aromatase gene *cyp19a1a* decreased only in some but not all transgenic females (Supplemental Fig. S6 H). Furthermore, the *gdf6Y* expression of both XX and XY transgenic animals was much higher than in the non-transgenic males due to the ubiquitous expression (Supplemental Fig. S6 I).

Hatching the mosaic *gdf6Y* transgenic animals, we observed that the hatchlings failed to straighten their body axes and were not able to swim normally (Supplemental Fig. S6 J). This detrimental phenotype was reminiscent of the homozygous loss of function (Fig. 2 E; Supplemental Fig. S2 H), suggesting that tight spatiotemporal regulation of the *gdf6* alleles is necessary for proper development in *N. furzeri*.

### Gdf6Y-responsive genes are differentially expressed between female and male *N. furzeri*

To identify Gdf6Y target genes, we transfected cell lines with a *gdf6Y* expression plasmid or one out of two control plasmids and analyzed their transcriptomes after 24 h. As a model for the somatic cells, which express *gdf6Y* in the killifish’s testes, we used murine Sertoli-like TM4 cells (Mather 1980). The transcriptome of TM4 cells transfected with the *gdf6Y* expression plasmid was compared with the transcriptome of TM4 cells either expressing the *gdf6Y* mutant variant *gdf6Y^del9^*or carrying an empty vector (EV). Overlaps of the DEGs from both comparisons resulted in 20 up- and 22 downregulated genes in the presence of *gdf6Y* (Fig. 5 A; Supplemental Fig. S7 A). An overrepresentation analysis revealed TGF-β signaling as the most enriched pathway among those 42 DEGs (Fig. 5 B). Out of the 7 genes assigned to this pathway, only *Tgfbr2* was downregulated, while *Id1*, *Id2*, *Id3*, *Smad6*, *Smad7*, and *Smad9* were upregulated (Fig. 5 A). With *Id1*, *Id2*, and *Id3* being known downstream effectors of BMPs (Hollnagel et al. 1999; Yang et al. 2013), this suggests that these genes are regulated as an immediate response to Gdf6Y-signaling. To verify the Gdf6Y-responsive genes that we had identified, we performed the same experiment in human HeLa cells (Fig. 5 C; Supplemental Fig. S7 B). Here, 6 commonly upregulated orthologous genes were identified, namely *Lxn*, *Id1*, *Id2*, *Id3*, and the inhibitory (I-) Smad-genes *Smad6* and *Smad7*, of which the latter 5 were previously assigned to the TGF-β signaling pathway (Fig. 5 A). The single commonly downregulated orthologous gene was *Zfp36l2*, which encodes a zinc finger protein that binds to AU-rich elements (AREs) in the 3’ untranslated regions (UTRs) of mRNAs promoting their degradation (Hudson et al. 2004). Expression analysis in accordingly treated human HEK293 cells confirmed the regulation of all these orthologous genes but *ID2* in the presence of *gdf6Y*, verifying them as general Gdf6Y-response genes (Supplemental Fig. S7 C). Next, we assessed their potential role in *N. furzeri* sex determination by examining the sex-specific expression of their orthologs in different tissues and developmental stages utilizing the previously published transcriptome data from Reichwald et al. (2015). While no reads had been mapped for the orthologs of *Id2* and *Lxn*, we found that *id1* was upregulated in male embryos at 10 dpf and *zfp36l2* was downregulated in testes of 3-month-old animals, corresponding to the *gdf6Y*-dependent regulation observed in the cell culture-based approach (Fig 5 D; Supplemental Fig. S7 D). Furthermore, the male-specific downregulation of *zfp36l2* in the gonad was reversed in phenofemales carrying either the *gdf6Y* mutant variant *gdf6Y^del9^*(Fig 5 E) or the frameshift mutation *gdf6Y^del113^* (Supplemental Fig. S7 E). These data suggest that the identified Gdf6Y-response genes, especially *id1*, and *zfp36l2*, are involved in *N. furzeri* sex determination.

**Figure 5.**
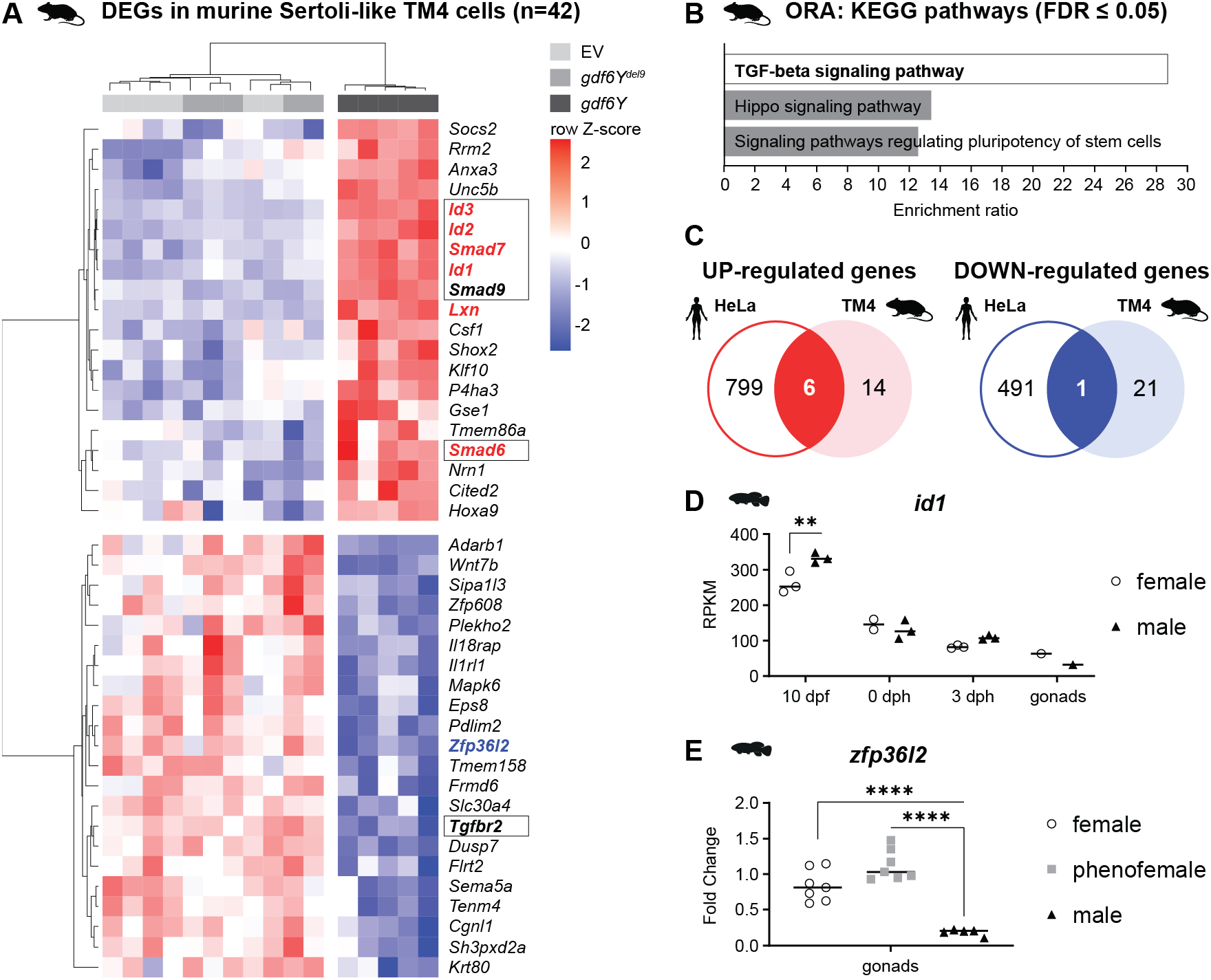
Gdf6Y leads to the specific regulation of certain genes independent of the targeted cell’s identity. *(A*) Heatmap of DEGs between murine Sertoli-like TM4 cells transfected with a *gdf6Y* expression plasmid and TM4 cells transfected with either a *gdf6Y* mutant variant (*gdf6Y^del9^*) expression plasmid or an empty vector (EV). *(B)* Overrepresentation analysis (ORA) of KEGG pathways within the 42 DEGs identified in TM4 cells. Genes of the “TGF-beta signaling pathway” are boxed and highlighted in bold letters in *(A)*. FDR – false discovery rate. *(C)* Overlap of the up-or downregulated DEGs identified in TM4 cells with those identified in human HeLa cells. The common genes are highlighted in red (upregulated genes) or blue (the downregulated gene) in *(A)*. *(D)* The expression of *id1* derived from RNA-Seq data in whole embryos at 10 dpf, embryo trunks at 0 or 3 dph, and gonads at 3 months of age in female and male *N. furzeri*. 2-way ANOVA and Šídák’s multiple comparisons test between females and males. (**) *P*<0.01. *(E)* RT-qPCR analysis of the *zfp36l2* expression in *N. furzeri* gonads of 3 months old females, males, and phenofemales carrying the *gdf6Y^del9^* mutation. 1-way ANOVA and Dunnett’s multiple comparisons test with males, (****) *P*<0.0001.

## DISCUSSION

Employing CRISPR/Cas9-mediated gene inactivation and overexpression obtaining full XY male-to-female and partial XX female-to-male sex reversals, respectively, we demonstrate that *gdf6Y* is necessary and sufficient for sex determination in *N. furzeri*. Among the identified *gdf6Y* mutations causing a male-to-female sex reversal, we found a 9 bp in-frame deletion within the prodomain (*gdf6Y^del9^*), which leads to the loss of amino acids 199 to 201. These amino acids are highly conserved among vertebrate *gdf6* orthologs and a heterozygous point mutation altering one of them leads to microphthalmia in humans (Asai-Coakwell et al. 2009). While not directly affecting the mature ligand, these residues might be important for prodomain folding and the release of the ligand from the proprotein complex, which is required for signaling. All other isolated *gdf6Y* mutants carried frameshift mutations leading to premature termination in the latter half of the prodomain. Similar frameshift mutations in *gdf6b* also led to male-to-female sex reversal in the Pachón cavefish *Astyanax mexicanus* (Imarazene et al. 2021), which harbors two *gdf6* paralogs, *gdf6a*, and *gdf6b*. While the SD gene in *A. mexicanus* arose from an additional *gdf6b* duplicate on a supernumerary B chromosome, *gdf6Y* and conversely *gdf6X* emerged through allelic diversification from an ancient *gdf6a* as no *gdf6b* ortholog is present in the available *N. furzeri* genome sequences (Reichwald et al. 2015; Valenzano et al. 2015).

While *gdf6Y* compensates for the loss of *gdf6X*, it also adopted the SD function in *N. furzeri* and possibly its sister species *N. kadleci,* which shares *N. furzeri*’s sex chromosomes (Stundlova et al. 2022). Three possible scenarios of how *gdf6Y*’s SD function evolved are functional alterations to the amino acid sequence, changes in the spatiotemporal gene expression, or a combination of both. The positive selection of *gdf6Y* in the SDR and the conservation of non-synonymous SNPs among strains support the theory of an allelic neofunctionalization (Reichwald et al. 2015). Structural modeling suggested that the four most C-terminal amino acid substitutions in Gdf6Y affect dimerization or ligand-receptor binding (Reichwald et al. 2015). We found that Smad2 activation by Gdf6Y was significantly increased compared to Gdf6X. Interestingly, a substitution of a highly conserved tyrosine for asparagine in human GDF6 was reported to decrease its sensitivity for the inhibitor Noggin (Wang et al. 2016). In *N. furzeri*, this residue is surrounded by the last two substitutions between Gdf6X and Gdf6Y, suggesting decreased inhibitor sensitivity as an explanation for Gdf6Y’s enhanced signaling capability. On the other hand, we confirmed the expression of *gdf6Y* but not *gdf6X* in adult *N. furzeri* testes, implying a differential regulation of the genes. We investigated DNA methylation as a potential cause of the differential expression of *N. furzeri*’s *gdf6* alleles in testes. While X-chromosomal hypermethylation of the SDR is crucial for sex determination in the channel catfish *Ictalurus punctatus* (Wang et al. 2022), the promoters of both *gdf6Y* and *gdf6X* are hypomethylated in gonads. The hypermethylation of the *sybu* promoter on the Y chromosome on the other hand indicates transcriptional silencing of *N. furzeri*’s Y-chromosome outside of the SD locus, suggesting an explanation for the lack of *rnf19a* expression in YY* embryos. The intergenic region between *gdf6X/gdf6Y* and *sybu* was highly methylated on both sex chromosomes. Therefore, the two Y-exclusive TEs in this region might not harbor *gdf6Y*-specific upstream enhancers, as enhancer activity would require a low or at most intermediate DNA methylation (Sharifi-Zarchi et al. 2017). In other fish species, TEs contribute sex-specific positive and negative regulatory elements that are crucial to SD gene function (Herpin et al. 2010; Herpin et al. 2021). Given that TEs accumulate on fish sex chromosomes during their early evolution (Chalopin et al. 2015) and contribute to the suppression of meiotic recombination (reviewed in Kent et al. (2017)), they could also cause an aberrant expression of non-SD genes in the SDR e.g., the Y-dependent expression of *pctp* in *N. furzeri* embryos. However, the Y-chromosomal effects on the expression of *rnf19a* and *pctp* might be GRZ-specific, as SDRs of other strains differ in size with the smallest containing only one gene, which is *gdf6Y* (Reichwald et al. 2015). Even allelic SNPs in *sybu* are not sex-linked in all strains (Reichwald et al. 2015). Therefore, the severe phenotype of YY* embryos would likely not occur in strains with smaller SDRs. Conversely, regulatory elements, causing the testis-specific expression of *gdf6Y*, should be located within the minimal SDR, which also contains a gene desert downstream of the gene itself. This well-conserved gene desert harbors most cis-regulatory elements controlling *GDF6* expression in other vertebrates (Mortlock et al. 2003; Portnoy et al. 2005; Pregizer and Mortlock 2009; Reed and Mortlock 2010; Indjeian et al. 2016; Bademci et al. 2020; Mortlock et al. 2022). Therefore, changes in or the acquisition of new Y-chromosomal enhancers downstream of *gdf6Y* are potential causes of its sex-specific expression. Whether protein or expression changes evolved first and which event constituted *gdf6Y* as the SD gene are open questions, but both probably played a role in its fixation.

Vertebrate sex determination is believed to be initiated by the sexual fate decision of the somatic cells of the bipotential gonadal primordium, which in turn control female or male gametogenesis. In a trade-off between male and female regulatory networks involving *Dmrt1* on the male and *Foxl2* on the female side, the equilibrium tips to one side by varying SD factors in different species (Herpin and Schartl 2015). In *N. furzeri*, *gdf6Y* is the male SD gene and is expressed in testicular somatic cells. In a cell culture-based approach murine Sertoli-like cells were used to identify Gdf6Y targets. In a complementary approach, Wang et al. (2022) analyzed gene expression in response to BMP signaling inhibition in human testis cells from 7-week-old fetuses, which is one to two weeks past the arrival of the migrating GCs in the genital ridge and, thereby, shortly after sex determination. As in our data, *ID1/2/3*, *SMAD6/7/9*, and *UNC5B* were upregulated in a BMP-dependent manner. However, only *UNC5B* was Sertoli cell-specific, while the other genes were regulated in all somatic and GC populations, showing the omnipresence of BMP signaling in the developing testis. This is consistent with our observation that *Id1/2/3* and *Smad6/7* are expressed in response to Gdf6Y, independent of cell type. *Arapaima gigas* provided evidence for the involvement of *ID* genes in sex determination in teleost species with a Y-chromosomal *id2b* duplicate being its SD gene (Adolfi et al. 2021). In *N. furzeri*, *id3* is positively selected like *gdf6Y*, suggesting a potential co-evolution (Reichwald et al. 2015). However, we found *id1* upregulated in male *N. furzeri* embryos at 10 dpf, indicating a role of Gdf6Y in paracrine signaling to the GCs during sex determination or sexual development. In mammals, *N. furzeri*, medaka, and zebrafish, female GCs enter meiosis during early gonadal development, while male GCs do not until puberty. Regarding the determination of the GCs’ sexual fate, *foxl2l* was identified as a GC-intrinsic feminizing factor in medaka, where the gene’s inactivation leads to a GC-specific female-to-male sex reversal and the development of sperm in ovaries that failed to initiate oogenesis at the regular developmental time point (Nishimura et al. 2015). While *foxl2l* inactivation in tilapia (*Oreochromis niloticus*) leads to a similar phenotype, it leads to a full female-to-male sex reversal in zebrafish (Dai et al. 2021; Liu et al. 2022). In *N. furzeri*, *foxl2l* is also involved in female GC sex determination, as shown by the masculinized expression of marker genes in mosaic *foxl2l* inactivated females and phenofemales at 0 dph. In medaka, male and female GCs initially express *foxl2l* at the onset of sexual differentiation of the gonads. While its expression is lost in male GCs, the mitotically active subpopulation of female GCs expresses *foxl2l* until entering meiosis throughout life (Nishimura et al. 2015; Nishimura and Tanaka 2016; Tanaka 2016). The role of *foxl2l-*expressing cells as pre-meiotic oocyte progenitors was confirmed in the zebrafish ovary (Liu et al. 2022). In *N. furzeri*, we also detected *foxl2l* expression in female GCs but not oocytes during ovarian development at 0 dph and in adulthood. In contrast, we rarely found *foxl2l* expressing GCs in developing testes at 0 dph and never in adult testes, supporting that its female GC SD function is conserved in *N. furzeri*. As the gonadal somatic cells direct GC fate during sex determination, Gdf6Y might be the somatic cell signal that initiates male GC fate and suppresses *foxl2l* expression. Therefore, *foxl2l* should be directly or indirectly downregulated by Gdf6Y-responsive genes like *id1*, which shows a male-specific expression at 10 dpf. In human fetal gonads, expression of TGF-β signaling pathway genes including *ID1/2/3* is enriched in both female and male GCs (Li et al. 2017). This observed TGF-β signaling was attributed to AMH from Sertoli cells in males and BMP2 from granulosa cells in females (Li et al. 2017). We detected *amh* expression in both types of somatic supporting cells in male and female *N. furzeri*. While AMH signals via the phosphorylation of the typical BMP-activated Smads 1, 5, and 9 (di Clemente et al. 2003; Orvis et al. 2008), Gdf6Y led to the phosphorylation of Smad9, Smad3, and Smad2 in an enhanced manner compared to Gdf6X. Interestingly, Smad2 is important in male GC sex determination in mice, where it is probably activated by the TGF-β family members activin and nodal (Wu et al. 2015). In *N. furzeri*, Gdf6Y could replace or interfere with this signaling. We also identified one downregulated gene in response to Gdf6Y: *zfp36l2*. It encodes an mRNA decay activator and is expressed in developing oocytes, where it is involved in meiotic cell cycle progression and oocyte maturation (Belloc and Mendez 2008; Ball et al. 2014; Ball et al. 2017; Sha et al. 2018), explaining its expression in *N. furzeri* female and phenofemale ovaries compared to testes. Zfp36l2 also controls epigenetic changes in the oocyte by regulating transcripts of histone and DNA demethylases (Chousal et al. 2018). While this takes place after sex determination, *zfp36l2* is female GC-specific and could influence gene expression on the transcriptional or indirectly the epigenetic level during GC sex determination. Overall, our data suggest that *gdf6Y* determines the male sex by suppressing female ovarian development. A similar mechanism was shown for the transcription factor SdY, which directly interacts with Foxl2 to suppress female differentiation in the gonadal somatic cells of *O. mykiss* (Bertho et al. 2018). However, while Gdf6Y could signal to both germ and somatic cells, our data suggests that it suppresses female differentiation of GCs instead of somatic cell.

## DATA AVAILABILITY

The RNA and bisulfite sequencing data discussed in this publication and applied in-house scripts are available on request.

## AUTHOR CONTRIBUTIONS

Conceptualization, A.R., C.E.; methodology, A.R.; software, R.S., P.K.; Formal Analysis, A.R., R.G., P.K., A.P.; investigation, A.R., H.M., M.T., M.K., R.G., C.A., N.H., A.H.; data curationA.R., H.M., M.T., M.K., R.G., C.A., P.K., M.G., A.P.; writing – original draft, A.R.; writing – review & editing, A.R., C.E.; visualization, A.R.; supervision, C.E.; funding acquisition, C.E.

## ACKNOWLEDGEMENTS

We thank Christina Ebert, Sabine Matz, Sabine Landmann-Weinsheimer, Silke Foerste, Peter Singer, Joseph Kölbel, and Gabriele Günther for their excellent technical support, and Hanna Reuter for her bioinformatical expertise. We are very grateful to members of the Core Facilities Next Generation Sequencing (Ivonne Goerlich, Cornelia Luge, and Martin Bens), Life Science Computing, and Imaging for their contributions. We also thank the members of FLI’s fish facility, most notably Simone Dunkel, Martin Neumann, Johannes Wilfert, Marcus Schmidt, Uta Naumann, and Beate Hoppe. This project was supported by a grant (EN 280/15–1) from the Deutsche Forschungsgemeinschaft to CE. The FLI is a member of the Leibniz Association and is financially supported by the Federal Government of Germany and the State of Thuringia.

## MATERIAL AND METHODS

### Experimental Models

#### Fish experiments

All animals were maintained in the fish facility of the Leibniz Institute on Aging – Fritz Lipmann Institute Jena according to the German Animal Welfare Law. All experiments were covered by animal experiment licenses 03-005/12, 03-044/16, and FLI-18-020 approved by the Thuringian authorities (Thüringer Landesamt für Verbraucherschutz).

All experiments were performed in the *N. furzeri* wild-type strain GRZ (Jubb 1971) or genetically modified lines in this strain background (Supplemental Tab. S1) except for the *in situ* hybridization in Supplemental Fig. S3 G, where MZM0403 was used instead (Terzibasi et al. 2008). Fish were single-housed at 26 °C in a light:dark cycle of 12 h each. Adult fish were once daily fed *ad libitum* with red mosquito larvae. Juveniles up to 5 weeks post-hatching were fed with artemia twice daily. To obtain fertilized GRZ oocytes for injections, breeding groups of five (one male, four females) to ten fish (two males, eight females) were set up in 20 to 40 l tanks, respectively. To obtain oocytes from phenofemales, breeding groups of one male and up to four phenofemales were set up in 8.5 l tanks. Sandboxes for egg deposition were removed for two days and placed back into the tank 2 h before injection. Eggs were collected with a sieve and used for microinjections. For line maintenance, eggs were collected weekly, put on coconut coir plates, and incubated at 29 °C to develop until 0 dph. For egg storage up to 1 year, eggs were incubated at 25°C to enter diapause and moved to 29 °C for 1 to 2 weeks before hatching.

Dechorionation of embryos and animals at 0 dph was performed as described previously (Richter et al. 2022). Phenotypes of embryos and larvae up to 2 dph were documented using reflective or transmission light with the Zeiss SteREO Discovery.V8 equipped with a Zeiss AxioCam MRc and a Volpi dual gooseneck pole mount light or the Zeiss Axio Zoom.V16 equipped with a Zeiss AxioCam HRc and an HXP 200 C light source, respectively. Phenotypes of adult animals were documented with a Canon IXUS 160.

#### Cell culture

TM4, HeLa, and HEK293 cells were cultivated in Dulbecco’s Modified Eagle Medium (DMEM; TM4, HeLa: Gibco™ #41966 – high glucose, pyruvate; HEK293: Gibco™ #31966 – high glucose, GlutaMAX™ Supplement, pyruvate) with 10 % FBS at 37 °C and 5 % CO_2_. For transfections, 0.5 × 10^6^ HeLa and TM4 cells or 4 × 10^5^ HEK293 cells per well were seeded into 6-or 12-Well plates, respectively. 24 h later, the cells were transfected with 2.5 µg or 1 µg of plasmid DNA (pcDgdf6Y, pcDgdf6Ydel9, or pcDNA3.1-HA) per well of a 6- or 12-Well plate, respectively, using Lipofectamine 3000 (Thermo Fisher Scientific) and incubated for 24 h at 37 °C and 5 % CO_2_ before RNA isolation.

Medaka fibroblast cells were cultured at 28°C and 5% CO2 in DMEM supplemented with high glucose (4.5 g/L), 4 mM glutamine, and 15% FBS as previously described (Thoma et al. 2011). For transfection, cells were grown to 80% confluency in six-well plates and subsequently transfected using either Lipofectamine (Thermo Fisher Scientific) or FuGENE (Roche) reagents as described by the manufacturers.

### Method Details

#### Tol2 transgenesis and CRISPR/Cas9 mutant generation

The plasmid pDTactb2:gdf6Y, which contains the *D. rerio actb2*-promoter driven *gdf6Y* expression cassette, was generated by combining the Tol2kit (Kwan et al. 2007) vectors #395 (pDestTol2CG2) and #299 (p5E-*bactin2*) with pMEgdf6Y and p3Egdf6YpA in a Gateway LR Clonase II (Thermo Fisher Scientific) reaction according to the manufacturer’s instructions. To generate plasmids pMEgdf6Y and p3Egdf6YpA, the *gdf6Y* genomic sequence from start- to stop-codon (including the intron) or the 965 bp sequence immediately downstream of *gdf6Y*, respectively, were amplified using Phusion High-Fidelity Polymerase (Thermo Fisher Scientific) according to the manufacturer’s instructions. The applied oligonucleotides added the respective *att*-sites to recombine the obtained amplicons into pDONR221 (Thermo Fisher Scientific) or pDONRP2R-P3 (Tol2kit #220) for the generation of pMEgdf6Y or p3Egdf6YpA, respectively, using Gateway BP Clonase II (Thermo Fisher Scientific) according to the manufacturer’s instructions. *Tol2*- and *EGFP*-mRNA were *in vitro* transcribed as described previously (Hartmann and Englert 2012) and quality-controlled by RNA agarose gel electrophoresis. The injection solution contained 20 ng/µl of pDTactb2:gdf6Y, 10 ng/µl of *Tol2*-mRNA, 30 ng/µl of *GFP*-mRNA, and 0.1 % phenol red.

To generate mutants with CRISPR/Cas9, sgRNAs were designed based on previously published X and Y chromosomal sequences (*gdf6Y-* and *gdf6X*-inactivation; Reichwald et al. (2015)) or by utilizing CHOPCHOP v3 (*foxl2l*-inactivation; Labun et al. (2019)). Templates for the *in vitro* transcription of sgRNAs were generated either by subcloning annealed oligonucleotides containing the target sequences into pDR274 (Addgene, plasmid #42250) via *BsaI* as described previously (*gdf6Y-*inactivation; Krug et al. (2023)) or by PCR (*gdf6X*-and *foxl2l*-inactivation). For the PCR, complementary oligonucleotides containing the T7 promoter, the 20 bp sgRNA target sequence, and the sgRNA backbone were applied as template-DNA at a concentration of 0.002 µM each and amplified using Phusion High-Fidelity Polymerase in 1X HF-Buffer (Thermo Fisher Scientific) with an additional primer pair (T7F and pDRrvs) according to the manufacturer’s instructions. The respective oligonucleotides are listed in Supplemental Tab. S2. The approximately 300 bp *DraI* restriction fragments (*gdf6Y-*inactivation) or the 124 bp amplicons (*gdf6X*- and *foxl2l*-inactivation) were purified from an agarose gel or the PCR, respectively, using the NucleoSpin Gel and PCR Clean-up Mini kit (Macherey-Nagel) according to the manufacturer’s instructions. Up to 1 µg purified DNA was *in vitro* transcribed overnight at 37°C with the MAXIscript T7 Transcription Kit (Thermo Fisher Scientific). The synthesized sgRNAs were precipitated from the reaction using Ammonium acetate Stop Solution as described for the mMessage mMachine Kit protocol (Thermo Fisher Scientific). *Cas9*-mRNA was *in vitro* transcribed overnight at 20°C from 400-500 ng *XbaI*-linearized pT3TS-nCas9n (Addgene, plasmid #46757) using the mMessage mMachine T3 Kit (Thermo Fisher Scientific) and purified with the MegaClear Kit (Thermo Fisher Scientific) according to the manufacturer’s instructions. All RNAs were quality-controlled by RNA agarose gel electrophoresis. Injection solutions contained 30 ng/µl of each sgRNA, 300 ng/µl of *Cas9*-mRNA, 30 ng/µl of *GFP*-mRNA, and 0.1 % phenol red. Microinjections into *N. furzeri* oocytes were performed as described previously by Hartmann and Englert (2012) (*gdf6Y*-inactivation) or Krug et al. (2023) (*gdf6Y*-transgenesis, *gdf6X*- and *foxl2l*-inactivation) applying the injection solutions described above.

#### Molecular sexing and genotyping of animals

Genomic DNA samples were derived from caudal fin biopsies of adult animals or tail tips from embryos and larvae as described previously by Krug et al. (2023) and Richter et al. (2022), respectively. For sexing and genotyping PCRs, DreamTaq DNA Polymerase (Thermo Fisher Scientific) was used according to the manufacturer’s instructions with an annealing temperature of 60°C, if not indicated otherwise. Sex chromosomes were determined by either a *gdf6Y*-specific PCR using oligonucleotides Nf_Gdf6m_fw4: 5’-GAGAAAGAGGAGCTGGTCGGG-3’ and Nf_Gdf6Y_rv4: 5’-GCGCTGCTGATGATGATGCC-3’ (Gonosome PCR in Supplemental Fig. S1 B) or as described previously (see Supplemental Fig. S2 B; Richter et al. (2022)). For the *gdf6Y*-specific PCR, an annealing temperature of 64°C was applied. As the *gdf6Y*-specific PCR flanks the target sites of sgRNA Y1 and Y2, it was also used for genotyping of *gdf6Y^del6, del8^*, and *gdf6Y^del9^*phenofemales yielding a 205 or 210 bp fragment, respectively, compared to a 219 bp fragment in males. *Gdf6Y^del113^* and *gdf6X^del113^*mutations were identified by applying oligonucleotides gdf6-gtF2: 5’-GGCCTCTGTGGGAGAATGTG-3’ and gdf6-gtR1: 5’-ACCGACCTCCGCGACTTG-3’ (Target locus PCR in Supplemental Fig. S1 B, C) indicating the mutant allele by a smaller fragment. Mutations in *foxl2l* were analyzed using oligonucleotides foxl2lt2F: 5’-GAGGTATGGATGCGGATAAGAA-3’ and foxl2lt3R: 5’-CTGATGCAGGTAGTAGTCGGTG-3’ flanking the sgRNA target sites. *Gdf6Y*-transgenic hatchlings and larvae were identified by the detection of EGFP in the heart by fluorescence microscopy and a transgene-specific 735 bp fragment by PCR applying oligonucleotides gdf6YinsituF1: 5’-CGGAGGGTAACCTGCTGC-3’ and M13F: 5’-GTAAAACGACGGCCAG-3’ at an annealing temperature of 55°C.

To determine mutation rates in *gdf6X/Y* in mosaic XY* animals, the *gdf6X/Y* target locus PCR was either performed as described above or using Phusion High-Fidelity Polymerase (Thermo Fisher Scientific) according to the manufacturer’s instructions. The obtained DreamTaq and Phusion amplified fragments were cloned into pCRII-TOPO using the TOPO TA Cloning Kit (Thermo Fisher Scientific) according to the manufacturer’s instructions or the pUC57-derivative pGGC (Geissler et al. 2011) via *SmaI* cut-ligation, respectively. Individual clones were Sanger sequenced with M13F in-house by the FLI Core Facility Next Generation Sequencing or externally by Eurofins Genomics TubeSeq Service. *Gdf6X/Y* target locus PCR amplicons from XX animals, injected with the *gdf6Y* sgRNAs Y1 and Y2, were purified from the reaction with NucleoSpin Gel and PCR Clean-up Mini kit (Macherey-Nagel) according to the manufacturer’s instructions and subjected to a T7 Endonuclease assay (T7EI) and Sanger sequencing with gdf6-gtF2 by Eurofins Genomics TubeSeq Service. For the T7EI assay, 200 ng of purified PCR product and 1 µl of NEB 2 Buffer (New England Biolabs) in 9.5 µl total volume were first denatured for 5 min at 95°C and then rehybridized by cooling the sample at a rate of 2°C/s to 85°C and a rate of 0.1°C to 25°C to obtain potential heteroduplex DNA. To detect potential heteroduplex DNA by cleavage, 0.5 µl of T7EI (New England Biolabs) were applied for 15 min at 37°C and the reaction was analyzed by agarose gel electrophoresis. Sanger sequences from individual clones and PCR products were analyzed with Geneious Prime 2019.2.3.

To determine *gdf6X/Y* mutation rates in mosaic animals, injected with the *gdf6X* sgRNAs X1 and X2, the *gdf6Y*-specific and the *gdf6X/Y* target locus PCR were performed on X*Y or X*Y and X*X* individuals, respectively. PCR products were Sanger sequenced with the forward primer of the respective PCR by Eurofins Genomics TubeSeq Service. Sequences of the *gdf6Y*-specific and the *gdf6X/Y* target locus PCR on X*X* animals were analyzed with Synthego Performance Analysis, ICE Analysis. 2019. v3.0. Synthego; [accessed between 03/05/2021 and 03/19/2021] by supplying the sgRNA Y2 or both *gdf6X*-sgRNA sequences, respectively. Sequences of the *gdf6X/Y* target locus PCR on X*Y animals were analyzed with Synthego Performance Analysis, ICE Analysis. 2019. v3.0. Synthego; [accessed between 03/05/2021 and 03/19/2021] by supplying both *gdf6X*-sgRNA sequences and the *gdf6Y*-sequence as a donor sequence to exclude the *gdf6Y* wild-type sequence from analysis of the *gdf6X* mutation rate. The result was evaluated by calculating the mutation rate as the percentage proportion of predicted indel sequences from the sum of indel and wild-type sequences.

PCR products from individual *foxl2l*-mosaic mutants were first examined for large indel mutations. Those samples without obvious indels were subjected to a T7EI assay as described above. T7EI-resistant samples were declared as not mutated, while PCR products from cleaved samples were Sanger sequenced with foxl2lt2F by Eurofins Genomics TubeSeq Service. Sequences were analyzed with Synthego Performance Analysis, ICE Analysis. 2019. v3.0. Synthego; [accessed on 11/17/2021 and 11/23/2021] by supplying the two *foxl2l*-sgRNA sequences and evaluated as described above.

#### Immunohistochemistry and RNAscope *in situ* hybridization

Dissected gonads and embryos were fixed in 4% paraformaldehyde/PBS overnight at 4°C, washed in phosphate-buffered saline, and paraffin-embedded. 5 μm sections were prepared with the Epredia HM 340 E electronic rotation microtome (Thermo Fisher Scientific) and transferred onto Superfrost Plus Slides (Thermo Fisher Scientific). The prepared sections were stained with Haematoxylin and Eosin (H&E) or used for RNAscope experiments. H&E stainings were imaged and processed with the AxioScan (Zeiss) and ZEN 2.6 (Blue Edition, Zeiss), respectively.

For RNAscope experiments, if not stated otherwise, buffers and equipment as well as probes were provided by Advanced Cell Diagnostics (Supplemental Tab. S3). The experiments were performed according to the manufacturer’s protocol (Advanced Cell Diagnostics, Document number: 323100-USM) except for the following changes: 60°C drying steps at the end of deparaffinization and target retrieval were prolonged from 5 to 30 min. Target retrieval and RNAscope Protease Plus treatment were performed for 15 and 30 min, respectively. After creating a barrier with the ImmEdge hydrophobic barrier pen, specimens were photobleached at 350-425 nm for 5 min with Axio Zoom.V16 or the SteREO Discovery.V8 (Zeiss) to reduce autofluorescence. All washing steps during hybridization and development were carried out three times for 5 min. The counterstaining with DAPI was either performed according to the manufacturer’s protocol or by applying and mounting the slides with ProLong Gold antifade reagent with DAPI (Thermo Fisher Scientific). For RNAscope samples, Z-stacks were recorded with a 63 x oil objective at the Axio Imager.Z1 with ApoTome.2 (Zeiss). Images were processed in ZEN 2.6 (Blue Edition, Zeiss) by combining all sections of Z-stack into a maximum-intensity projection.

#### Whole mount *in situ* hybridization

MZM0403 hatchlings at 0 dph were fixed in 4% paraformaldehyde/PBS and processed for whole mount *in situ* hybridization as described previously (Hauptmann and Gerster 1994). The digoxigenin-labeled antisense and sense *sybu* riboprobes were synthesized from a *SacII*-or *SacI*-linearized pGEM-T easy (Promega), which contained an 823 bp cDNA-fragment from *sybu*-mRNA (NCBI Reference Sequence: XM_015949383.2) amplified with oligonucleotides Nf_sybu_fw2: 5’-AGAGGACCATCAGCACCAAC-3’ and Nf_sybu_rv6: 5’-CAGGGACTCCAACTTCCT-3’, using the plasmids SP6 or T7 promoter, respectively. Prior probe hybridization, samples were bleached in 6 % hydrogen peroxide/PBST for 1 h at room temperature, followed by a 30 min Proteinase K-treatment. After probe detection, hatchlings were again fixed in 4% paraformaldehyde/PBS, taken through a methanol series, equilibrated, and oriented for imaging in a clearing solution (1/3 benzyl benzoate, 2/3 benzyl alcohol). Images were acquired using the Zeiss SteREO Discovery.V8 equipped with a Zeiss AxioCam MRc. Images were processed in ZEN 2.6 (Blue Edition, Zeiss). Sex was determined subsequently by sequencing amplicons generated with oligonucleotides scaffold795_4r: 5’-AAATCCTCTCCTGGCCTACC-3’ and scaffold795_3f: 5’-GGCTGGCGAGAGTCTTTCTA-3’ at genomic DNA isolated from the individual samples.

#### Bisulfite sequencing

Genomic DNA from testes or ovaries was isolated using the QIAamp DNA Mini Kit (QIAGEN) according to the protocol of the manufacturer. The deamination was performed with the EZ DNA Methylation-Gold Kit (Zymo Research) according to the manufacturer’s instructions. After deamination, samples were stored at -20°C. PCRs were performed to amplify the region of interest. The reaction mix was composed of 10 µl MyTaq HS Red Mix (Meridian Bioscience), 2 µl deaminated DNA and the respective oligonucleotides at a concentration of 0.5 µM each in 20 µl total volume, and amplified using the following program: initial denaturation at 95°C for 3 min, 30 cycles of denaturation at 95°C for 30 s, annealing at 55°C for 30s and elongation at 72°C for 1 min, followed by a final elongation at 72°C for 5 min. Amplicons were quality-controlled by agarose gel electrophoresis and distributed in equal amounts into 8 pools. The amplicons per pool (7 pools of 34 amplicons each, 1 pool of 27 amplicons) were individually tagged by one out of four 3 bp-tags (ATG, GCA, CAT, TGC), which were added to the oligonucleotides used for the amplification. The pools of amplicons were purified using Agencourt AMPure XP magnetic beads (Beckman Coulter) according to the manufacturer’s instruction. Amplicons pools were quality checked using the 2100 Bioanalyzer instrument in combination with the DNA 7500 assay (both Agilent Technologies). For each pool, an individually tagged library was prepared from 125 ng of input material using the TruSeq DNA PCR-Free kit (Illumina). Since pools of amplicons do not need to be fragmented, the protocol was started with the “Repair Ends” step. The quantification and quality check of libraries were done as described above. Libraries were pooled and sequenced on the MiSeq system. It ran in 151 cycle/paired-end mode (MiSeq reagent kit v2 - 300 cycles). Sequence information was converted to FASTQ format using bcl2fastq v2.20.0.422.

#### RNA isolation

To isolate RNA from *N. furzeri* gonads and transfected cells, the QIAGEN RNeasy Mini Kit was used. Killifish tissue was lysed in RLT buffer containing β-mercaptoethanol using the QIAGEN TissueLyzer II according to the manufacturer’s instructions. Cells were lysed directly in the wells using RLT buffer containing ß-mercaptoethanol. Cell samples were further homogenized using the QIAshredder Kit (QIAGEN). DNase digestion was performed on-column using the RNase-Free DNase Set (QIAGEN) as suggested in the RNeasy Mini Kit protocol. The RNA was eluted in 50 µl DEPC-treated H_2_O.

Total RNA from embryo and hatchling trunks was extracted by adding TRIzol reagent (Thermo Fisher Scientific) and homogenization with the QIAGEN TissueLyser II. For phase separation, the lysate was incubated for 5 min at room temperature followed by the addition of 0.2 volumes of chloroform relative to the starting amount of TRIzol reagent. The solution was shaken vigorously for at least 15 s, incubated for 3 min at room temperature, and centrifuged for 20 min at 12000 × g at 4°C. The aqueous phase was transferred into a fresh Tube and 1 volume of chloroform was added. Shaking, incubation at room temperature, centrifugation, and transfer of the aqueous phase into a fresh Tube were repeated. Alternatively, the lysate was transferred into Phasemaker tubes (Thermo Fisher Scientific), and the phase separation was performed in a single step according to the manufacturer’s instructions. For RNA precipitation, 0.5 μl GlycoBlue Coprecipitant (Thermo Scientific Scientific), 1.1 volumes of isopropanol, and 0.16 volumes of 2 M sodium acetate (pH 4.0) were added. After mixing and incubation for 10 min at room temperature, samples were centrifuged for 20 min at 12000 × g and 4°C. The supernatant was removed and 500 μl of 80 % ethanol were added to wash the pellet by vortexing and centrifugation at 7500 × g for 10 min at 4°C. The washing step was repeated. The pellet was air-dried for 10-15 min and dissolved in 15 to 30 μl of DEPC-treated H_2_O. For optimal dissolving of the pellet, the sample was incubated at 65°C for 5 min. For RT-qPCR of *foxl2l*, samples were cleaned-up and DNase digested with the QIAGEN RNeasy Mini Kit and the RNase-Free DNase Set (QIAGEN) according to the manufacturer’s instructions.

#### Reverse transcription-quantitative PCR (RT-qPCR)

For cDNA Synthesis up to 1 µg of total RNA was used. The reverse transcription was performed using the iScript cDNA Synthesis Kit (Bio-Rad) according to the protocol provided by the manufacturer. RT-qPCR was performed in 384 well plates using the CFX384 Real-Time system (Bio-Rad). Each 10 µl reaction contained 5 µl SYBR GreenER™ qPCR SuperMix Universal (Thermo Fisher Scientific), forward and reverse primer at a concentration of 200 nM each, and up to 5 ng/µl cDNA and was cycled according to the manufacturer’s instruction. Annealing temperatures were 60°C or 63°C for *N. furzeri* or human cell assays, respectively. Oligonucleotides and their applications are listed in Supplemental Tab. S4.

#### RNA sequencing

Sequencing of RNA samples was performed using Illumina’s next-generation sequencing methodology (Bentley et al. 2008). In detail, DNase digested total RNA from TM4, and HeLa cells was quantified, and quality checked using the Tapestation 4200 instrument in combination with an RNA ScreenTape (both Agilent Technologies). Libraries were prepared from 300 ng total RNA using NEBNext Ultra II Directional RNA Library Preparation Kit in combination with NEBNext Poly(A) mRNA Magnetic Isolation Module and NEBNext Multiplex Oligos for Illumina (Unique Dual Index UMI Adaptors RNA) following the manufacturer’s instructions (New England Biolabs). Deviating from the instructions, qPCR was performed for each library prior to final amplification to determine the optimal number of cycles (exponential phase) to avoid over-amplification of the libraries in the final amplification step. The quantification and quality check of libraries was done using the Bioanalyzer 2100 instrument and the D5000 kit (Agilent Technologies). Libraries were pooled and sequenced on a NovaSeq6000. The system ran in 101 cycle/single-end/standard loading workflow mode (S1 100 cycle kit v1.5). Sequence information was converted to FASTQ format using bcl2fastq v2.20.0.422.

#### Smad-activation luciferase reporter assay

The luciferase reporter assay to analyze Smad-activation mediated by the different *N. furzeri gdf6* alleles and *gdf6Y* mutant variants was performed as reported previously (Herpin et al. 2021). In short, medaka fibroblast cells in six-well plates were co-transfected with 400 ng of one out of four *gdf6*-expression plasmids (pcDgdf6X, pcDgdf6Y, pcDgdf6Ydel9, or pcDgdf6Yins25), 300 ng of one out of five expression plasmids for fusion proteins consisting of Smad activation and Gal4 DNA binding domains (Smad1-AD-GAL4-DBD, Smad2-ADGAL4-DBD, Smad3-AD-GAL4-DBD, Smad5-AD-GAL4-DBD, or Smad8-AD-GAL4-DBD), 300 ng of a firefly luciferase reporter plasmid under the control of a UAS sequence containing minimal promoter (UAS-luc), and 5 ng of a *Gaussia* luciferase expression plasmid for normalization (pCMV-Gluc) per well. After 24 h, cells were washed with 2× phosphate-buffered saline and lysed with 75 µL of passive lysis buffer (Promega), and then subjected to luciferase assay. Firefly luciferase activity was quantified using the Dual-Luciferase Reporter Assay System (Promega) and normalized against *Gaussia* luciferase activity.

### Quantification and statistical analysis

#### RT-qPCR quantification

RT-qPCR results were evaluated using the determined threshold cycles (Ct) and primer efficiencies (E) either by calculating the relative expression (R) of a gene of interest (goi) to a reference gene (ref) with the formula:

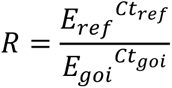

(Supplemental Fig. S3 A and S4 C) or the fold change (FC) of a goi to one or two reference genes (ref1, ref2) with the formula:

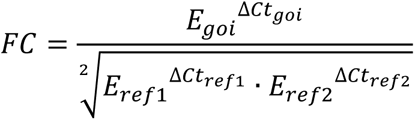

based on Hellemans et al. (2007) (all other RT-qPCR data).

#### Statistical analyses

Graphical presentation and statistical analyses of RT-qPCR data, retina layer measurements, mutation rates, luciferase assay results and comparisons of individual genes’ RPKM from the *N. furzeri* RNA-Seq data were realized using GraphPad Prism 9 and the indicated statistical test depending on the scientific question and data structure of the respective assay.

#### Bisulfite sequencing data analyses

For data analyses, two operating systems were used. All Python-based software pool_separator.py, methi.py) was controlled with a SunOS 5.10 system. To analyze their output, R studio was used on a Windows 10 operating system. Merging of the paired-end reads per amplicon pool was done using clc_overlap_reads (CLC-Workbench, QIAGEN). The Python in-house script pool_separator.py was used to separate the amplicons stored in the joined .fastq-files. To supply the script with the necessary information, a .csv-file including the semicolon-separated information (pool; 3 bp-tag; forward primer; reverse primer; sample name) for each line-separated amplicon was generated for each pool. The command to run the Python script is shown below with pool 1 as an example:

python3 pool_separator.py Ampliconpool1_jointed_test.fq pool1comp.csv

The script creates a folder for each amplicon, which contains all .fastq-files of each individual sample. The in-house script methi.py was used to calculate the CpG methylation rates. This script separates X and Y chromosomal reads and detects the chromosome-specific methylation rates for all CpGs of each X and Y separated amplicon. The analysis of each amplicon by the script requires the forward and reverse PCR oligonucleotide sequences, the X- and Y-specific amplicon length, the tag length, the position of all CpGs within the amplicon (comma-separated), a location of a non-CpG cytosine as well as 9 to 10 bp X (p1) and Y (p2) signature sequences (9 – 10 bp). The information is stored in a .txt-file as shown below:

forward_primer:
reverse_primer:
expected_length_p1:
expected_length_p2:
tag_length:
3 cpg_pos_p1:
cpg_pos_p2:
cx_pos_p1:
cx_pos_p2:
p1:
p2:

The command to run the software methyi.py is shown below:

python3 software/methi.py -d fastq_files/ -i sample_name.txt -s

The cytosine, which is not in a CpG context, is needed used by methyi.py to estimate the conversion rate of the deamination process and to correct the methylation rate. The program uses the frequency (H) of the non-CpG (CpX, X: cytosine or thymine) to calculate the conversion rate (c) as follows:

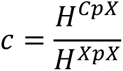

The median (c_all_) over all CpG methylation rates of the amplicons from all samples was calculated in R Studio as follows:

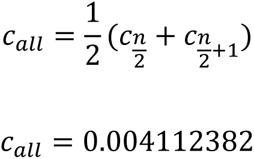

Subsequently, the c_all_ was used to calculate the corrected methylation rate (CpG_corr_) with a value from 0 to 1 for each CpG:

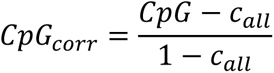

Alternatively, the methylation rate of a CpG was determined sample-wise with Geneious Prime 2019.2.3 by assembling the pool-separated reads to a bisulfite-conversed reference sequence of the respective amplicon and measuring the proportion of cytosines at the given position. The obtained methylation rate was corrected with the formula above.

The corrected methylation rates of all analyzed CpGs and samples were visualized using Morpheus (https://software.broadinstitute.org/morpheus).

#### RNA-Seq data analyses

The published RNA-Seq data (ENA: ERR879038 - ERR879056) from male and female *N. furzeri* at different developmental stages (Reichwald et al. 2015) was filtered with sga (parameters: preprocess -q 30 -m 50 --dust; https://github.com/jts/sga). Quality-passed reads were mapped to the *N. furzeri* reference genome using STAR STAR (parameters: --alignIntronMax 100000; Dobin et al. (2013)). DEGs (adjusted p-value < 0.05, Log_2_(Fold Change) > 1) between females and males of each individual developmental stage were derived using DESeq2 (Love et al. 2014). DEGs were visualized in volcano plots generated with VolcaNoseR (Goedhart and Luijsterburg (2020); https://huygens.science.uva.nl/VolcaNoseR2/).

Analysis of RNA-Seq data of murine TM4 and human HeLa cells. The names of the RNA-Seq reads were extended by their unique molecular identifier (UMI) with UMI-tools’ extract command version 1.1.1 (Smith et al. 2017). These modified reads were aligned with STAR 2.7.10a (parameters: --alignIntronMax 100000, --outSJfilterReads Unique, --outSAMmultNmax 1, --outFilterMismatchNoverLmax 0.04) (Dobin et al. 2013) to the *M. musculus* (GRCm39) and *H. sapiens* (GRCh38) reference genomes, respectively, with the Ensembl genome annotation release 107. PCR duplicates were removed from the alignment results using UMI-tools’s dedup command. For each gene, reads that map uniquely to one genomic position were counted with FeatureCounts 2.0.3 (multi-mapping or multi-overlapping reads were discarded, stranded mode was set to “–s 2”) (Liao et al. 2014). The pairwise comparisons of *gdf6Y*-expressing samples with either *gdf6Y^del9^*-expressing or empty vector samples were analyzed for differential gene expression. Therefore, DEGs were determined with R 4.1.3 using the package DESeq2 1.34.0 (Love et al. 2014). Only genes that have at least one read count in any of the analyzed samples of a particular comparison were subjected to DESeq2. For each gene, the p-value was calculated using the Wald significance test. The resulting p-values were adjusted for multiple testing with the Benjamini & Hochberg correction (FDR). The log2 fold change (log2FC) values were shrunk with the DESeq2 function lfcShrink(type=“apeglm”) to control for variance of log2FC estimates for genes with low read counts (Zhu et al. 2019). Genes with an adjusted p-value < 0.05 are considered DEGs. Selected DEGs were visualized by calculating Z-scores (formula: 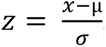, x = individual value, µ = arithmetic mean of all x per gene, σ = standard deviation of all x per gene) from DESeq2-normalized counts and creating a heatmap of those using Morpheus (https://software.broadinstitute.org/morpheus). The overlap analysis of the up-or downregulated DEGs identified in TM4 cells with those identified in the same way in human HeLa cells was based on gene symbols and followed by an ortholog verification in Ensembl release 109.

Overrepresentation analyses of KEGG pathways were performed using WebGestalt (WEB-based GEne SeT AnaLysis Toolkit; Liao et al. (2019); https://www.webgestalt.org/). For the *N. furzeri* RNA-Seq data, gene symbols from *D. rerio* orthologs assigned to the genebuild_v1.150922 *N. furzeri* annotation (Reichwald et al. 2015; Kelmer Sacramento et al. 2020) were used to search for enriched KEGG pathways in a *D. rerio* organismal background. Therefore, female or male DEGs at the different developmental stages were compared to a reference list of all genes identified in the whole data set. For the TM4 RNA-Seq data, gene symbols of the 42 selected DEGs were compared against the *M. musculus* genome to identify enriched KEGG pathways in that genetic background. Parameters for the enrichment analyses were a minimum number of 5 IDs in a category, a maximum number of 2000 IDs in a category and the Benjamini & Hochberg correction for multiple testing.

---

**Supplemental Figure S1.**
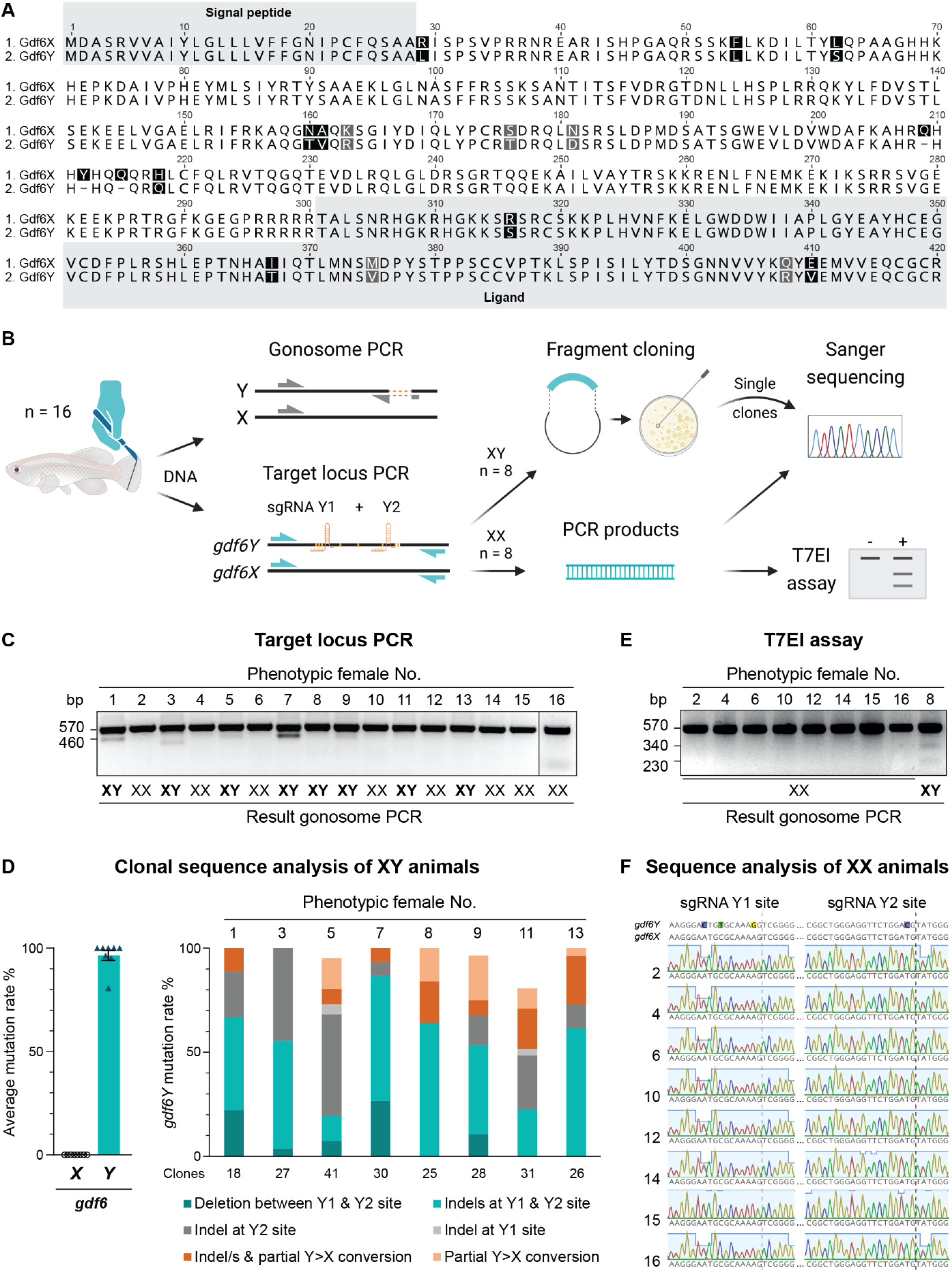
Comparison of Gdf6X and Gdf6Y and analysis of mosaic *gdf6Y* mutants. *(A*) Alignment of 1. Gdf6X and 2. Gdf6Y amino acid sequences (one-letter code). Sequence differences are highlighted in gray for similar and black for dissimilar amino acids. *(B)* Schematic of *gdf6Y* mutant analysis (Created with BioRender.com). T7EI – T7 endonuclease I. *(C)* Ethidium bromide-stained agarose gel picture of the sgRNA Y1 and Y2 target locus PCR on fin tissue lysates from the 16 phenotypically female CRISPR/Cas9-mutated animals. The sex chromosomes of each individual, as derived from the gonosome PCR, are indicated below the samples. *(D)* Result of the clonal sequence analysis of the 8 CRISPR/Cas9-mutated XY animals. Left: Average mutation rates in *gdf6X* and *gdf6Y* with SEM and individual values. Right: *Gdf6Y* mutation rates in the individual animal samples are separated into the different observed mutation types. Apart from insertion and deletion (indel) events at either or both sgRNA target sites, partial gene conversion between *gdf6X* and *gdf6Y* was observed probably due to homology-directed repair of DNA double-strand breaks. *(E)* Ethidium bromide-stained agarose gel picture of the T7EI assay on target locus PCR products from the 8 CRISPR/Cas9-mutated XX animals. No mutations were detected compared to the CRISPR/Cas9-mutated XY animal sample. *(F)* Partial sequence alignments with chromatograms of target locus PCR products from the 8 CRISPR/Cas9-mutated XX animals. No double peaks or decreased sequence qualities (light blue columns) were detected in chromatograms at the putative Cas9 cleavage sites mediated by sgRNA Y1 and Y2 (dashed lines).

**Supplemental Figure S2.**
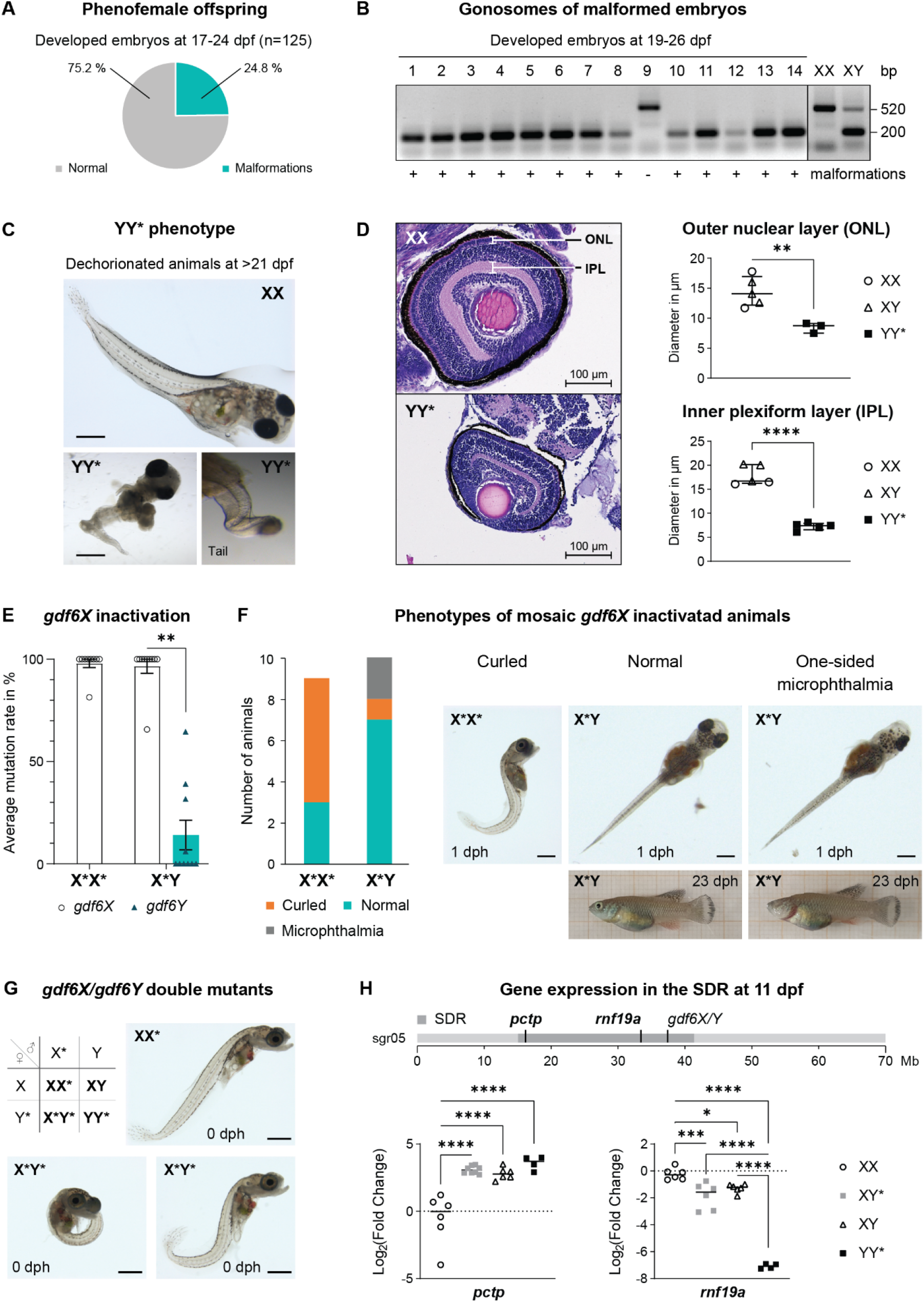
YY* embryo and *gdf6X* mutant phenotypes. *(A*) One quarter of a phenofemale’s (*gdf6Y^del9^*) offspring develops malformations. *(B)* Ethidium bromide-stained agarose gel picture of a molecular sexing assay utilizing an amplicon length polymorphism between the X and Y chromosomes. In malformed embryos, only the Y-chromosomal fragment was detected indicating their identity as YY* embryos (*gdf6Y^del6, del8^*). *(C)* Dechorionated YY* embryos (*gdf6Y^del113^*) have unpigmented and malformed tails as well as smaller eyes than normally developing e.g., XX, animals at 0 dph. *(D)* Left: H&E staining of normally developed (XX) and underdeveloped YY* (*gdf6Y^del6, del8^*) eyes at 0 dph or 19-26 dpf, respectively. Right: Quantification of diameters (median with interquartile range) of the outer nuclear layer (ONL) and the inner plexiform layer (IPL) of the retinas of normally developed and YY* (*gdf6Y^del6, del8^*) animals. The ONL consists of the photoreceptor cell bodies and the IPL contains the synapses between the bipolar cell axons and the dendrites of the ganglion and amacrine cells. Layer diameter was measured at three different positions per sample using ZEN 2.6 (Blue Edition, Zeiss) and averaged. Statistical testing by a two-tailed, unpaired *t*-test. *(E)* Result of the sequence analysis with the Synthego ICE tool (v3.0) of the 19 *gdf6X*-mosaic animals created with CRISPR/Cas9. Statistical testing by a Wilcoxon matched-pairs signed rank test corrected for multiple testing with the Holm-Šídák method. *(F)* Distribution according to gonosomes and representation of the phenotypes of *gdf6X*-mosaic animals. Predominantly females (X*X*) had a detrimental curled phenotype, while one-sided microphthalmia was present only in two males (X*Y). *(G) Gdf6X*/*gdf6Y* double mutant hatchlings (X*Y*; n = 30) have a detrimental curled phenotype (two manifestations shown), which prevents regular swimming behavior, while siblings (XX*, XY; n = 59) are normal in this regard. XX* shown for comparison. *(C, F, G)* Scale bars: 500 µm. *(H)* Schematic of the chromosomal location and expression of *pctp* and *rnf19a* in trunks of XX, XY*, XY, and YY* embryos at 11 dpf. SDR – sex-determining region. Sgr05 – synteny group 5. Statistical testing by a 1-way ANOVA and Tukey’s post hoc test. *(D, E, H)* (**) *P*<0.01, (***) *P*<0.001, (****) *P*<0.0001.

**Supplemental Figure S3.**
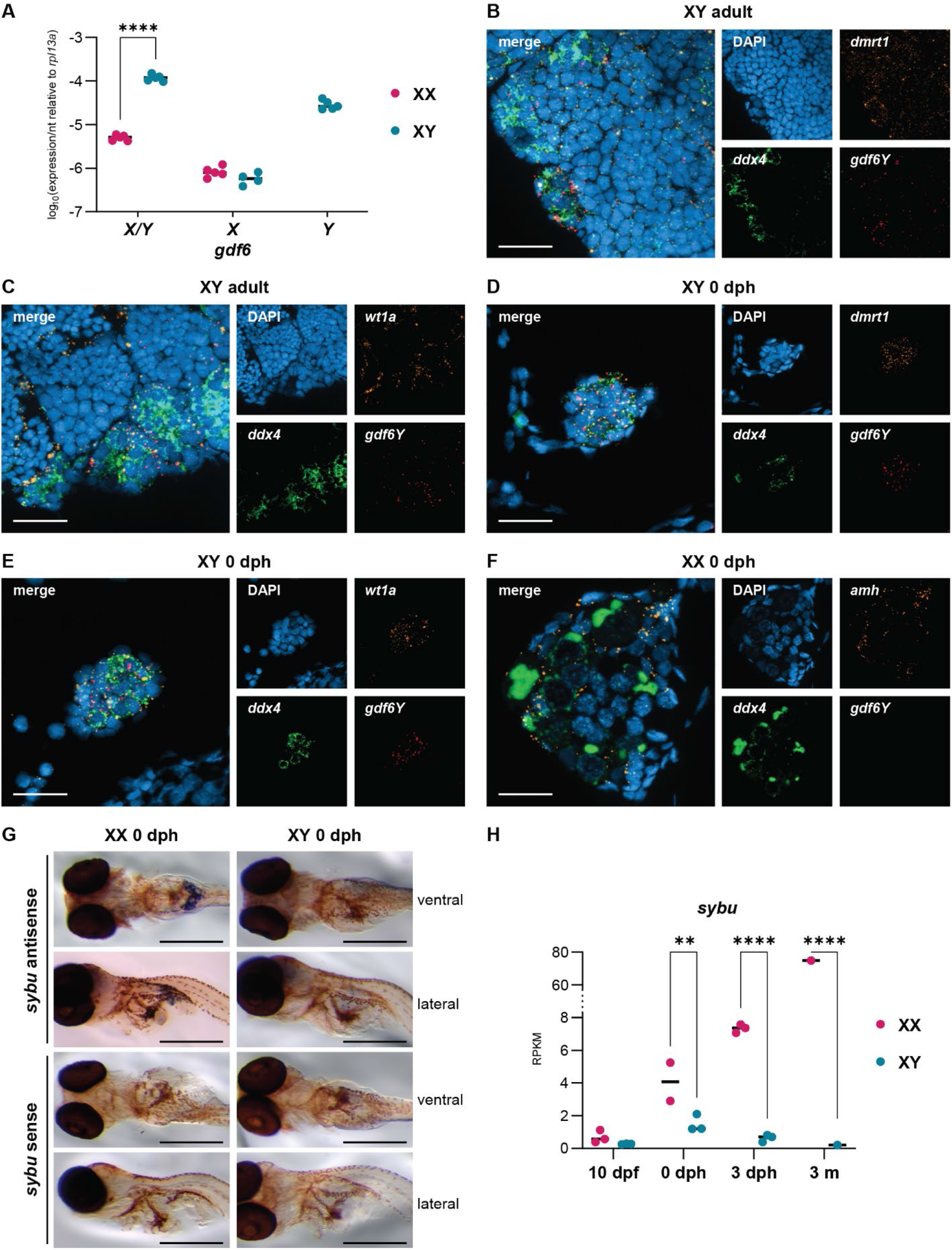
*Gdf6Y* expression in *N. furzeri*. *(A)* Expression of *gdf6X* and *gdf6Y* together and separately relative to *rpl13a* in gonads of adult females (XX) and males (XY). *(B, C, D, E, F)* Transcripts of *gdf6Y* (red) and the GC marker *ddx4* (green) as well as one somatic cell marker were detected. Scale bars: 20 µm. The somatic cell markers *dmrt1 (B, D)* and *wt1a (C, E)* were detected in the testes of adult *(B, C)* and 0 dph animals *(D, E). (F)* The somatic cell marker *amh* (orange) was detected in a 0 dph ovary. *(G)* Whole mount *in situ* hybridization of MZM0403 hatchlings with *sybu*-mRNA antisense (females: n = 5, males: n = 3) and sense probes (n = 8; definitive females: n = 3, definitive males: n = 3). *(H)* Expression analysis of *sybu* by RNA-Seq in the whole embryo at 10 dpf, the trunk at 0 or 3 dph, and the gonad at 3 months (m) of age in XX and XY animals. RPKM – Reads per kb of transcript per Million mapped reads. *(A, H)* Statistical testing by a 2-way ANOVA and Šidák’s post hoc test between XX and XY, (**) *P*<0.01, (****) *P*<0.0001.

**Supplemental Figure S4.**
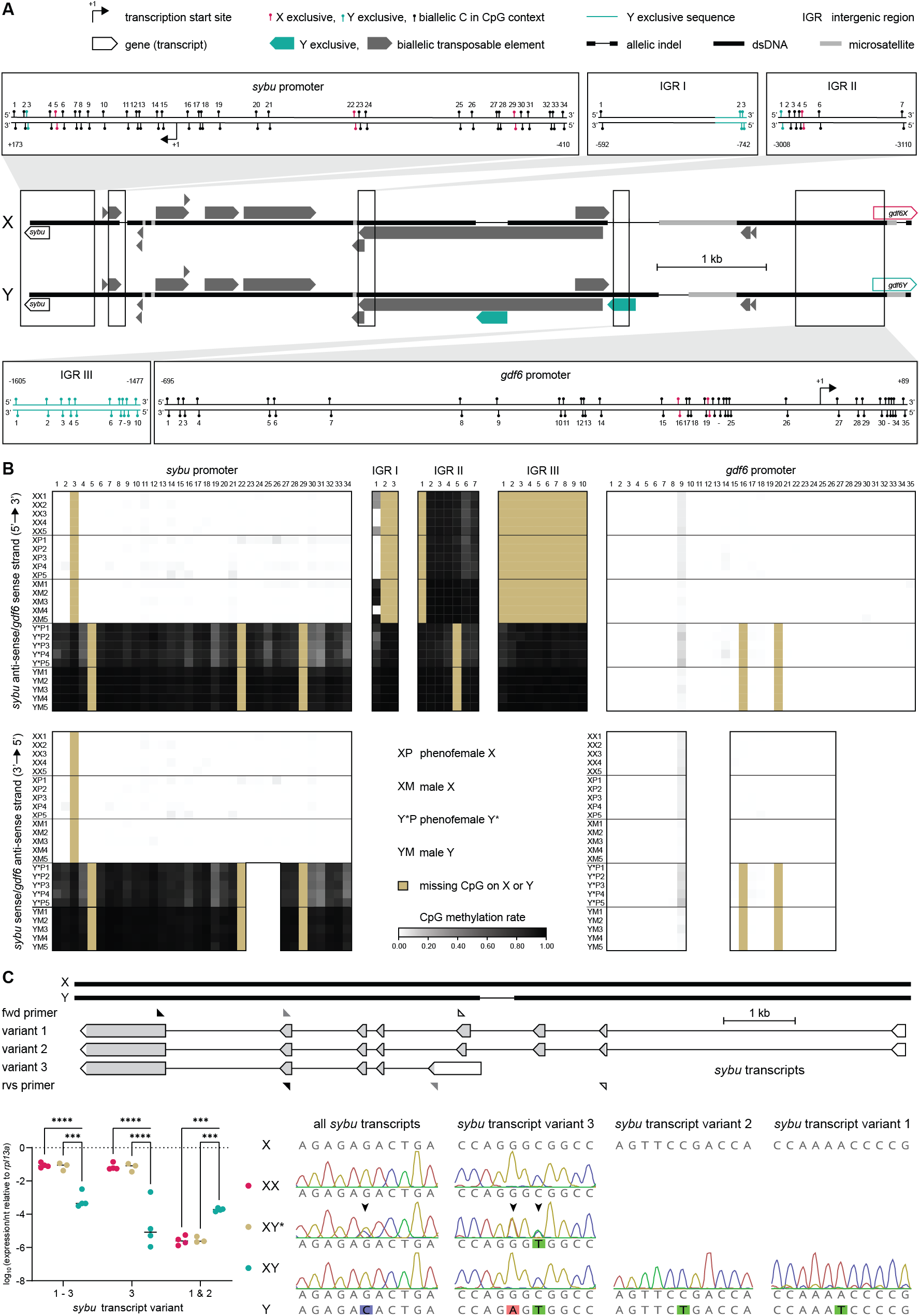
DNA methylation of the genomic region between *sybu* and *gdf6X/Y* on the sex chromosomes in gonads. *(A)* Schematic of the analyzed region of the X and Y chromosome containing Y exclusive TEs (*gdf6Y*-distal TE: NfRep007207 of unknown identity; *gdf6Y*-proximal TE: partial retrotransposon NfRep000041 of LINE/RTE-BovB type; source: https://nfingb.leibniz-fli.de). Insets show the CpG positions of the analyzed subregions. *(B)* Heatmaps of CpG methylation rates on the X and Y chromosomes in the gonads of the analyzed individuals (n=5). For some locations, the *sybu* sense strand (bottom panels) was analyzed in addition to the *gdf6X/Y* sense strand (upper panels). *(C)* Schematic and sex-specific expression of three different *sybu* transcripts in the gonads of females (XX), phenofemales (XY*), and males (XY). Exons and transcript-specific primer pairs are indicated with triangles (black – transcript variants 1-3, gray – variant 3, white – variant 1 and 2). *Sybu* cDNA sequences indicate an X-exclusive transcription in males, while expression from the Y chromosome is reactivated in phenofemales. Statistical testing by a 2-way ANOVA and Tukey’s post hoc test, (***) *P*<0.001, (****) *P*<0.0001.

**Supplemental Figure S5.**
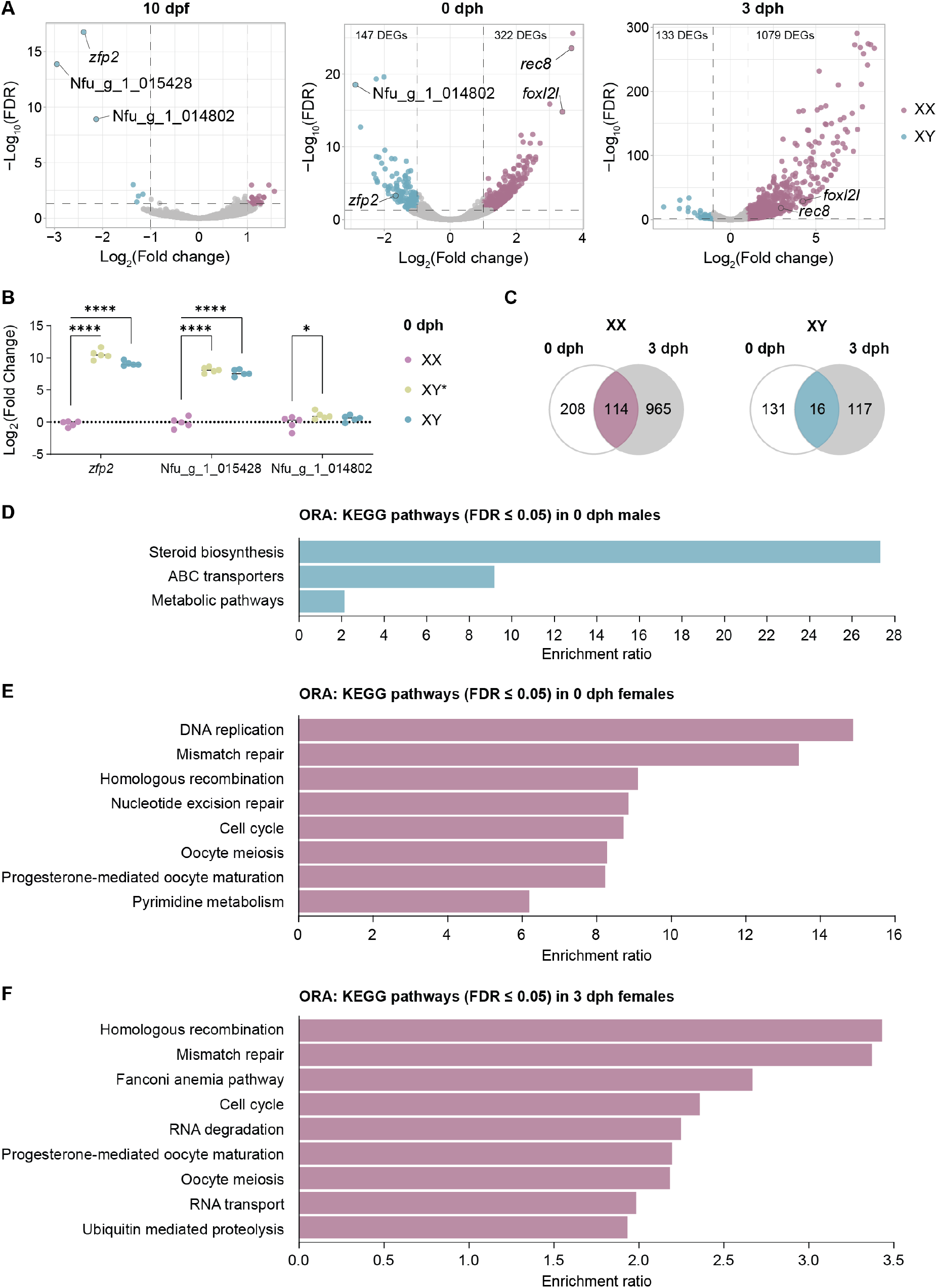
RNA-Seq results show that ovarian development starts before 0 dph in *N. furzeri*. *(A)* Volcano plots of DEGs between XX and XY animals in the whole embryo at 10 dpf and in trunk parts at 0 and 3 dph as derived from a respective previously published data set (XX at 0 dph n=2, others n=3; Reichwald et al. 2015). *(B)* Expression of the three top DEGs at 10 dpf in trunks of XX, XY*, and XY animals at 0 dph. Statistical testing by a 1-way ANOVA and Dunnett’s multiple comparisons test with XX animals, (*) *P*<0.05, (****) *P*<0.0001. *(C)* Overlap of female and male DEGs between 0 and 3 dph. *(D, E, F)* Overrepresentation analysis (ORA) of KEGG pathways within the male-*(D)* and female-specific *(E)* DEGs at 0 dph as well as the female-specific *(F)* DEGs at 3 dph.

**Supplemental Figure S6.**
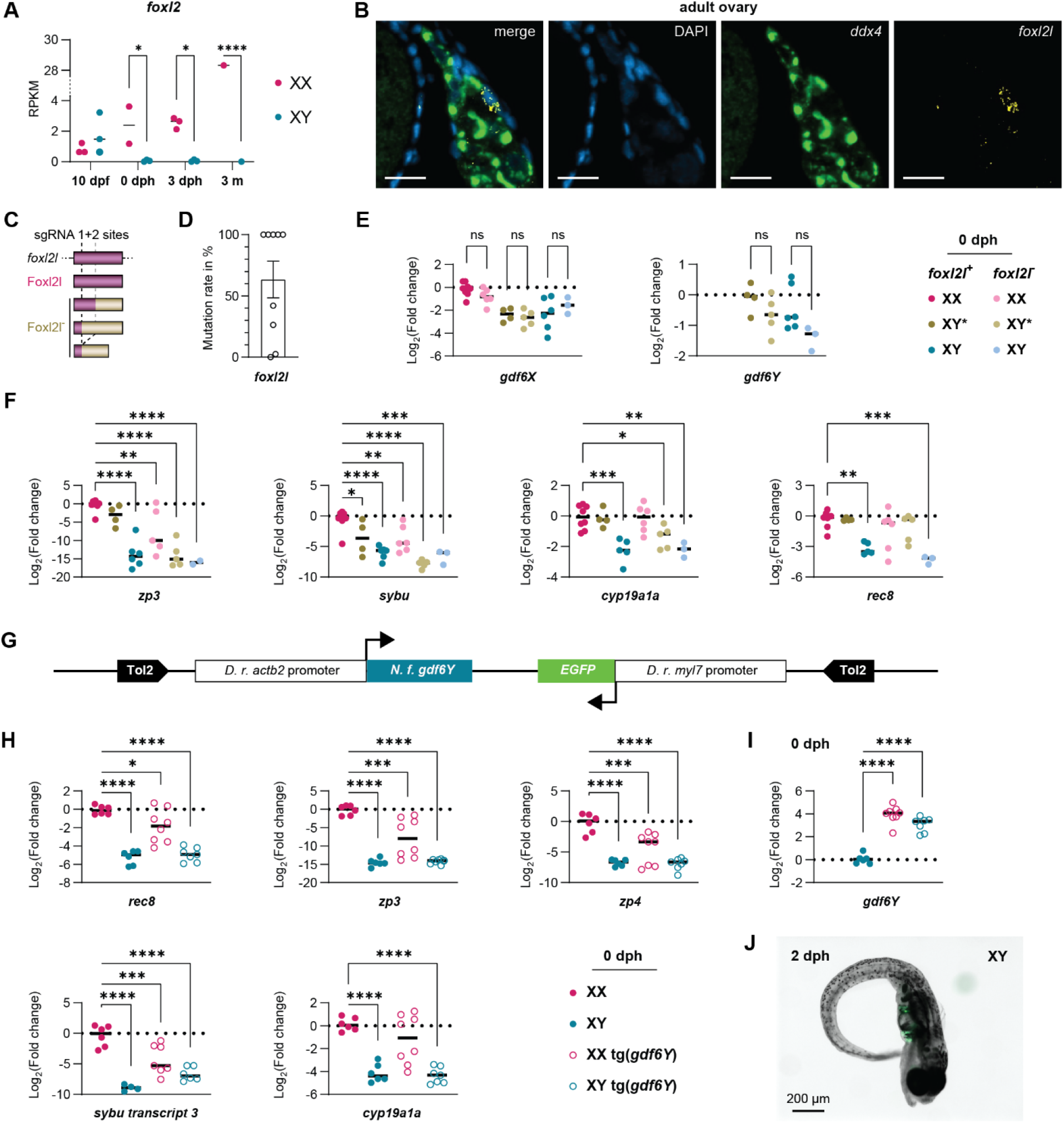
Expression of oogenesis-associated genes in mosaic *foxl2l*-mutants and *gdf6Y*-transgenic animals. *(A*) Expression analysis of *foxl2* by RNA-Seq in the whole embryo at 10 dpf, the trunk at 0 or 3 dph, and the gonad at 3 months (m) of age in XX and XY animals. 2-way ANOVA and Šidák’s post hoc test. *(B)* Ovarian expression at 5 weeks of age of *foxl2* (yellow) and the GC marker *ddx4* (green). Nuclei are stained with 4′,6-Diamidin-2-phenylindol (DAPI, blue). Scale bar: 20 µm. *(C)* Frameshifts induced by one or both *foxl2l* sgRNAs lead to truncated Foxl2l-proteins (Foxl2l^-^). *(D)* Result of sequence analyses with the Synthego ICE tool (v3.0) of the 9 *foxl2l*-mosaic animals created with CRISPR/Cas9. *(E)* Expression of *gdf6X* and *gdf6Y* in trunks of females (XX), phenofemales (XY*), and males (XY) and their mosaic *foxl2l*^-^ counterparts at 0 dph. Statistical testing by a 1-way ANOVA and Holm-Šídák’s multiple comparisons tests. *(F)* Expression of female marker genes in trunks of females, phenofemales, and males and their mosaic *foxl2l*^-^ counterparts at 0 dph. *(G)* Schematic of Tol2-transgene to drive ubiquitous *gdf6Y-*expression with a *D. rerio* (*D. r*.) *actb2* promoter in *N. furzeri* (*N. f.*). Heart-specific *EGFP expression* by a *D. r. myl7* promoter is used as a selection marker. *(H)* Expression of female marker genes in trunks of females, males, and their mosaic *gdf6Y*-transgenic (tg(*gdf6Y*)) counterparts at 0 dph. *(F, H)* Statistical testing by a 1-way ANOVA and Dunnett’s multiple comparisons test with XX animals. *(I)* Expression of *gdf6Y* in trunks of females, males, and their mosaic *gdf6Y*-transgenic counterparts at 0 dph. Statistical testing by a 1-way ANOVA and Dunnett’s multiple comparisons test with XY animals. *(E, F, H, I)* (*) *P*<0.05, (**) *P*<0.01, (***) *P*<0.001, (****) *P*<0.0001. *(J)* Curled *gdf6Y*-transgenic XY-larva with heart-specific EGFP expression (green).

**Supplemental Figure S7.**
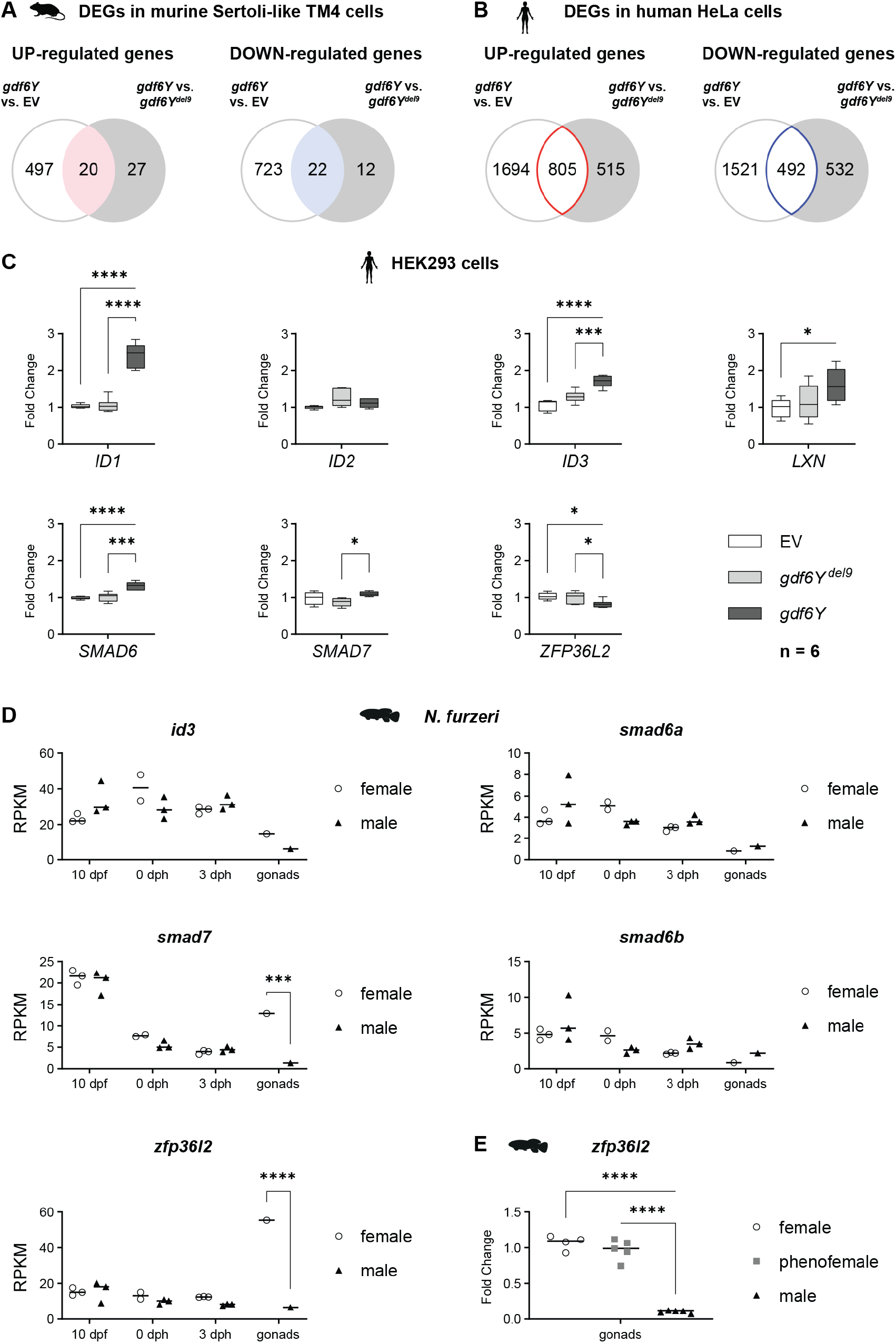
Gdf6Y-responsive genes in different cell lines and *N. furzeri*. *(A*) Overlap of the up-or downregulated DEGs between murine Sertoli-like TM4 cells transfected with either a *gdf6Y* or a *gdf6Y* mutant variant (*gdf6Y^del9^*) expression plasmid and TM4 cells transfected with either a *gdf6Y* expression plasmid or an empty vector (EV). *(B)* Overlap of the up-or downregulated DEGs between human HeLa cells transfected with either a *gdf6Y* or a *gdf6Y* mutant variant (*gdf6Y^del9^*) expression plasmid and TM4 cells transfected with either a *gdf6Y* expression plasmid or an empty vector (EV). *(C)* RT-qPCR analysis of the genes commonly up-or downregulated in TM4 and HeLa cells in human HEK293 cells transfected with an expression plasmid for *gdf6Y* or a *gdf6Y* mutant variant (*gdf6Y^del9^*) or an empty vector (EV). 1-way ANOVA and Dunnett’s multiple comparisons test with the *gdf6Y* transfected cells. *(D)* The expression of *id3*, *smad6a*, *smad6b*, *smad7*, and *zfp36l2* derived from RNA-Seq data in whole embryos at 10 dpf, embryo trunks at 0 or 3 dph and gonads at 3 months of age in female and male *N. furzeri*. 2-way ANOVA and Šídák’s multiple comparisons test between females and males. *(E)* RT-qPCR analysis of the *zfp36l2* expression in *N. furzeri* gonads of 3 months old females, males, and phenofemales carrying the frameshift mutation *gdf6Y^del113^*. 1-way ANOVA and Dunnett’s multiple comparisons test with males. *(C, D, E)* (*) *P*<0.05, (***) *P*<0.001, (****) *P*<0.0001.

## SUPPLEMENTAL TABLES

**Supplemental Table S1.**
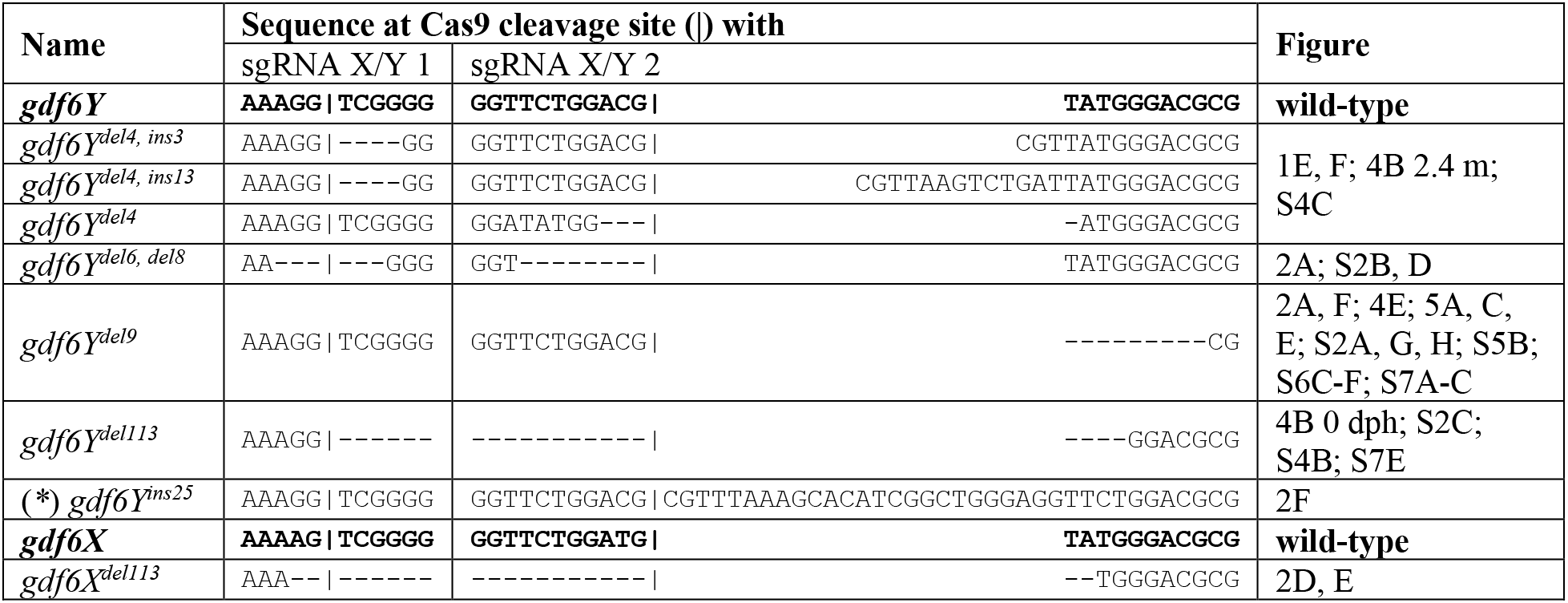
Mutations of *gdf6Y* and *gdf6X* used in this work. (*) Artificial mutation used in *in vitro* analysis.

**Supplemental Table S2.**
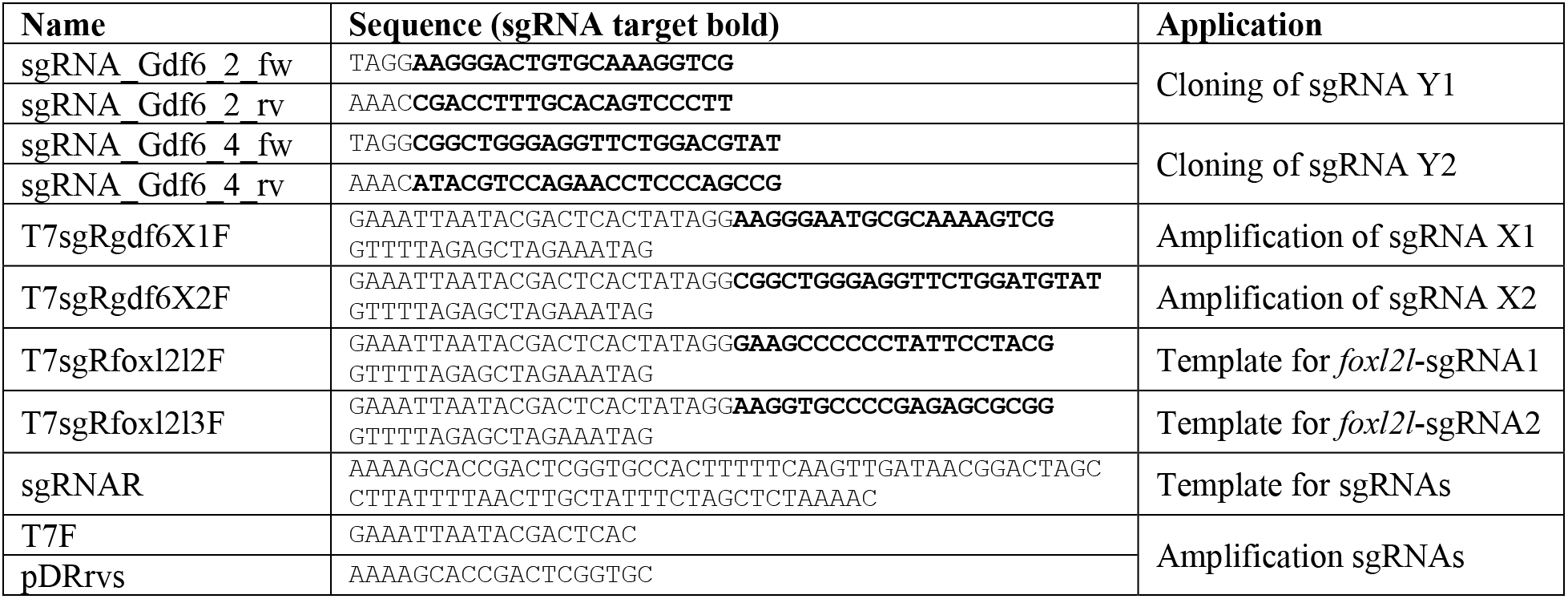
Oligonucleotides used for the synthesis of sgRNA *in vitro* transcription templates.

**Supplemental Table S3.** RNAscope probes purchased from Advanced Cell Diagnostics.

| Name | Targeted transcript<br>(NCBI Reference Sequence) | Catalogue-No. |
| --- | --- | --- |
| RNAscope® Probe - Nf-ddx4 | XM 015957842.1 (*ddx4*) | 563361 |
| RNAscope® Probe - Nf-amh-C2 | XM 015977279.1 (*amh*) | 830891-C2 |
| RNAscope® Probe - Nf-LOC107386104-C2 | XM 015960292.1 (*dmrt1*) | 830901-C2 |
| RNAscope® Probe - Nf-LOC107389222-C2 | XM 015965254.1 (*wt1a*) | 830871-C2 |
| RNAscope® Probe - Nf-LOC107390636-C2 | XM 015967458.1 (*foxl2l*) | 831201-C2 |
| RNAscope® Probe - Nf-gdf6y-C3 | *gdf6Y*-transcript (Reichwald et al. 2015) | 563381-C3 |

**Supplemental Table S4.**
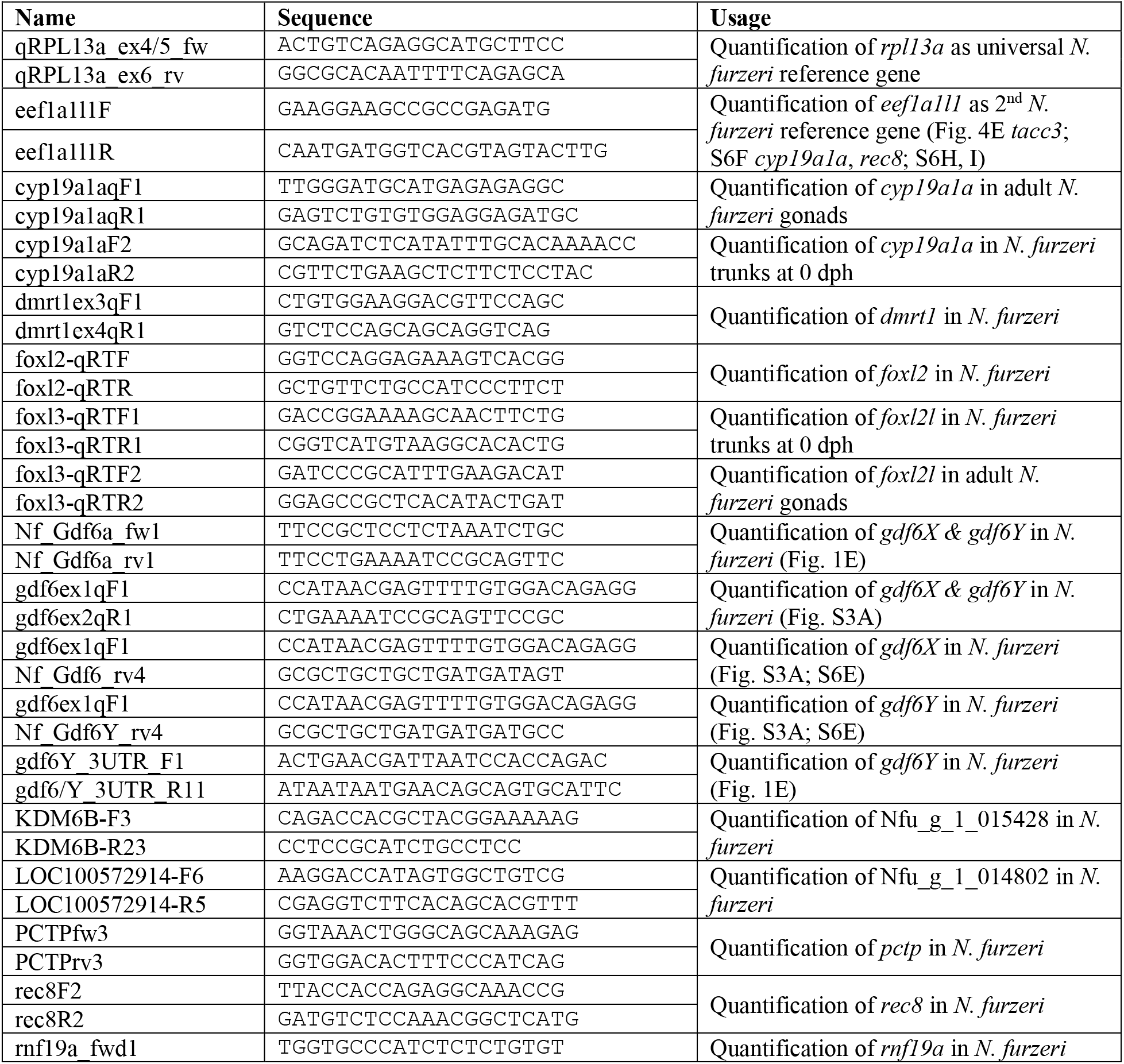
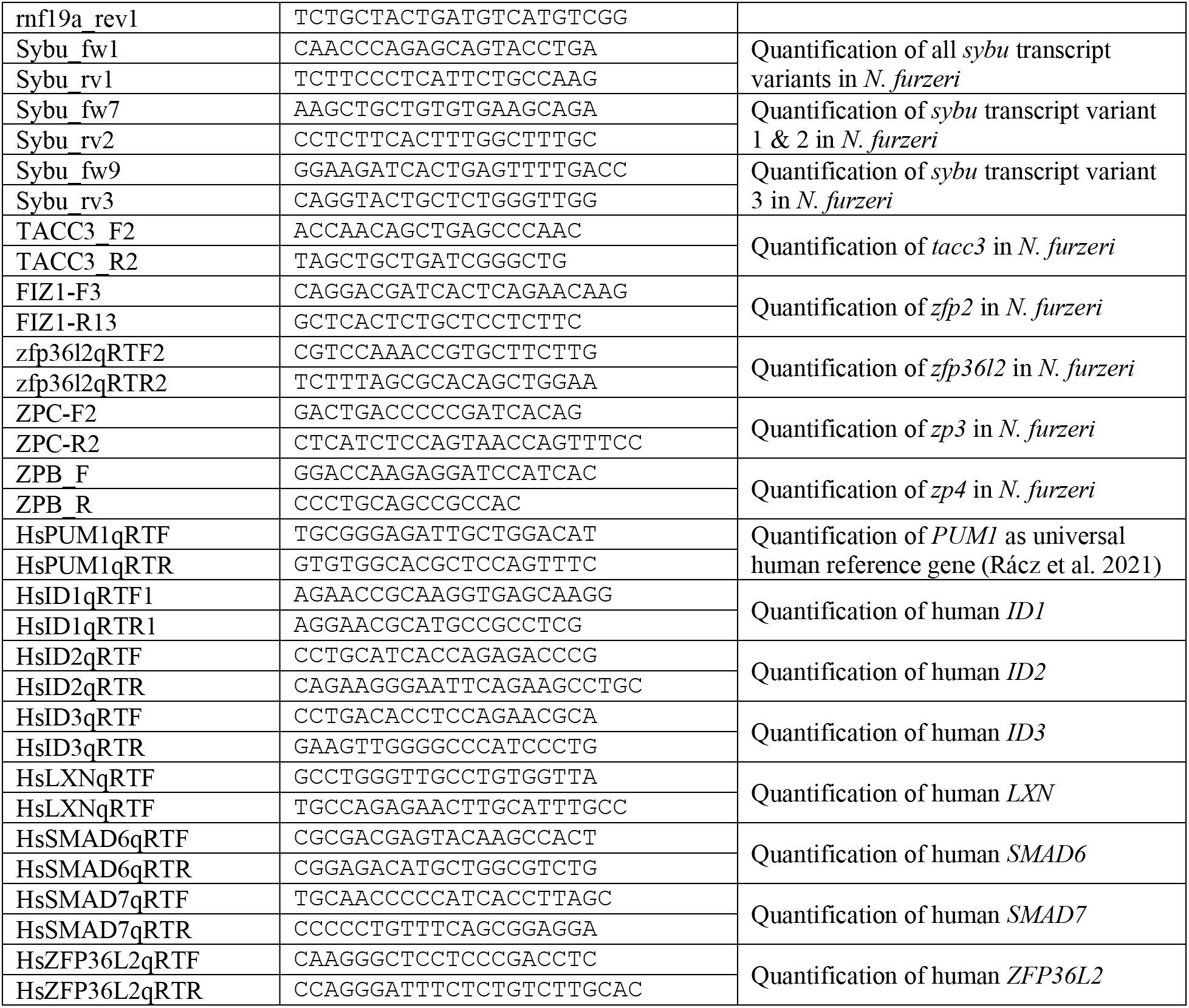
Oligonucleotides used for RT-qPCR.

## REFERENCES

1. Adolfi MC, Du K, Kneitz S, Cabau C, Zahm M, Klopp C, Feron R, Paixao RV, Varela ES, de Almeida FL et al. 2021. A duplicated copy of id2b is an unusual sex-determining candidate gene on the Y chromosome of arapaima (Arapaima gigas). Sci Rep 11: 21544.

2. Api M, Biondi P, Olivotto I, Terzibasi E, Cellerino A, Carnevali O. 2018. Effects of Parental Aging During Embryo Development and Adult Life: The Case of Nothobranchius furzeri. Zebrafish 15: 112–123.

3. Asai-Coakwell M, French CR, Berry KM, Ye M, Koss R, Somerville M, Mueller R, van Heyningen V, Waskiewicz AJ, Lehmann OJ. 2007. GDF6, a novel locus for a spectrum of ocular developmental anomalies. American journal of human genetics 80: 306–315.

4. Asai-Coakwell M, French CR, Ye M, Garcha K, Bigot K, Perera AG, Staehling-Hampton K, Mema SC, Chanda B, Mushegian A et al. 2009. Incomplete penetrance and phenotypic variability characterize Gdf6-attributable oculo-skeletal phenotypes. Human molecular genetics 18: 1110–1121.

5. Asai-Coakwell M, March L, Dai XH, Duval M, Lopez I, French CR, Famulski J, De Baere E, Francis PJ, Sundaresan P et al. 2013. Contribution of growth differentiation factor 6-dependent cell survival to early-onset retinal dystrophies. Human molecular genetics 22: 1432–1442.

6. Bademci G, Abad C, Cengiz FB, Seyhan S, Incesulu A, Guo S, Fitoz S, Atli EI, Gosstola NC, Demir S et al. 2020. Long-range cis-regulatory elements controlling GDF6 expression are essential for ear development. The Journal of clinical investigation 130: 4213–4217.

7. Ball CB, Rodriguez KF, Stumpo DJ, Ribeiro-Neto F, Korach KS, Blackshear PJ, Birnbaumer L, Ramos SB. 2014. The RNA-binding protein, ZFP36L2, influences ovulation and oocyte maturation. PloS one 9: e97324.

8. Ball CB, Solem AC, Meganck RM, Laederach A, Ramos SBV. 2017. Impact of RNA structure on ZFP36L2 interaction with luteinizing hormone receptor mRNA. RNA 23: 1209–1223.

9. Belloc E, Mendez R. 2008. A deadenylation negative feedback mechanism governs meiotic metaphase arrest. Nature 452: 1017–1021.

10. Bentley DR Balasubramanian S Swerdlow HP Smith GP Milton J Brown CG Hall KP Evers DJ Barnes CL Bignell HR, et al. 2008. Accurate whole human genome sequencing using reversible terminator chemistry. Nature 456: 53–59.

11. Bertho S, Herpin A, Branthonne A, Jouanno E, Yano A, Nicol B, Muller T, Pannetier M, Pailhoux E, Miwa M et al. 2018. The unusual rainbow trout sex determination gene hijacked the canonical vertebrate gonadal differentiation pathway. Proc Natl Acad Sci U S A 115: 12781–12786.

12. Chalopin D, Volff JN, Galiana D, Anderson JL, Schartl M. 2015. Transposable elements and early evolution of sex chromosomes in fish. Chromosome Res 23: 545–560.

13. Chousal J, Cho K, Ramaiah M, Skarbrevik D, Mora-Castilla S, Stumpo DJ, Lykke-Andersen J, Laurent LC, Blackshear PJ, Wilkinson MF et al. 2018. Chromatin Modification and Global Transcriptional Silencing in the Oocyte Mediated by the mRNA Decay Activator ZFP36L2. Dev Cell 44: 392–402 e397.

14. Colozza G, De Robertis EM. 2014. Maternal syntabulin is required for dorsal axis formation and is a germ plasm component in Xenopus. Differentiation 88: 17–26.

15. Crosier PS, Kalev-Zylinska ML, Hall CJ, Flores MV, Horsfield JA, Crosier KE. 2002. Pathways in blood and vessel development revealed through zebrafish genetics. The International journal of developmental biology 46: 493–502.

16. Dai S, Qi S, Wei X, Liu X, Li Y, Zhou X, Xiao H, Lu B, Wang D, Li M. 2021. Germline sexual fate is determined by the antagonistic action of dmrt1 and foxl3/foxl2 in tilapia. Development (Cambridge, England) 148.

17. den Hollander AI, Biyanwila J, Kovach P, Bardakjian T, Traboulsi EI, Ragge NK, Schneider A, Malicki J. 2010. Genetic defects of GDF6 in the zebrafish out of sight mutant and in human eye developmental anomalies. BMC genetics 11: 102.

18. di Clemente N, Josso N, Gouedard L, Belville C. 2003. Components of the anti-Mullerian hormone signaling pathway in gonads. Molecular and cellular endocrinology 211: 9–14.

19. Dobin A, Davis CA, Schlesinger F, Drenkow J, Zaleski C, Jha S, Batut P, Chaisson M, Gingeras TR. 2013. STAR: ultrafast universal RNA-seq aligner. Bioinformatics 29: 15–21.

20. Feron R, Zahm M, Cabau C, Klopp C, Roques C, Bouchez O, Eche C, Valiere S, Donnadieu C, Haffray P et al. 2020. Characterization of a Y-specific duplication/insertion of the anti-Mullerian hormone type II receptor gene based on a chromosome-scale genome assembly of yellow perch, Perca flavescens. Mol Ecol Resour 20: 531–543.

21. Geissler R, Scholze H, Hahn S, Streubel J, Bonas U, Behrens SE, Boch J. 2011. Transcriptional activators of human genes with programmable DNA-specificity. PloS one 6: e19509.

22. Goedhart J, Luijsterburg MS. 2020. VolcaNoseR is a web app for creating, exploring, labeling and sharing volcano plots. Sci Rep 10: 20560.

23. Gosse NJ, Baier H. 2009. An essential role for Radar (Gdf6a) in inducing dorsal fate in the zebrafish retina. Proc Natl Acad Sci U S A 106: 2236–2241.

24. Gramann AK, Venkatesan AM, Guerin M, Ceol CJ. 2019. Regulation of zebrafish melanocyte development by ligand-dependent BMP signaling. eLife 8.

25. Hall CJ, Flores MV, Davidson AJ, Crosier KE, Crosier PS. 2002. Radar is required for the establishment of vascular integrity in the zebrafish. Developmental biology 251: 105–117.

26. Hanel ML, Hensey C. 2006. Eye and neural defects associated with loss of GDF6. BMC developmental biology 6: 43.

27. Hartmann N, Englert C. 2012. A microinjection protocol for the generation of transgenic killifish (Species: Nothobranchius furzeri). Developmental dynamics : an official publication of the American Association of Anatomists 241: 1133–1141.

28. Hattori RS, Murai Y, Oura M, Masuda S, Majhi SK, Sakamoto T, Fernandino JI, Somoza GM, Yokota M, Strussmann CA. 2012. A Y-linked anti-Mullerian hormone duplication takes over a critical role in sex determination. Proc Natl Acad Sci U S A 109: 2955–2959.

29. Hauptmann G, Gerster T. 1994. Two-color whole-mount in situ hybridization to vertebrate and Drosophila embryos. Trends in genetics : TIG 10: 266.

30. Hellemans J, Mortier G, De Paepe A, Speleman F, Vandesompele J. 2007. qBase relative quantification framework and software for management and automated analysis of real-time quantitative PCR data. Genome biology 8: R19.

31. Herpin A, Braasch I, Kraeussling M, Schmidt C, Thoma EC, Nakamura S, Tanaka M, Schartl M. 2010. Transcriptional rewiring of the sex determining dmrt1 gene duplicate by transposable elements. PLoS genetics 6: e1000844.

32. Herpin A, Schartl M. 2015. Plasticity of gene-regulatory networks controlling sex determination: of masters, slaves, usual suspects, newcomers, and usurpators. EMBO reports 16: 1260–1274.

33. Herpin A, Schartl M, Depince A, Guiguen Y, Bobe J, Hua-Van A, Hayman ES, Octavera A, Yoshizaki G, Nichols KM et al. 2021. Allelic diversification after transposable element exaptation promoted gsdf as the master sex determining gene of sablefish. Genome research 31: 1366–1380.

34. Holborn MK, Einfeldt AL, Kess T, Duffy SJ, Messmer AM, Langille BL, Brachmann MK, Gauthier J, Bentzen P, Knutsen TM et al. 2022. Reference genome of lumpfish Cyclopterus lumpus Linnaeus provides evidence of male heterogametic sex determination through the AMH pathway. Mol Ecol Resour 22: 1427–1439.

35. Hollnagel A, Oehlmann V, Heymer J, Ruther U, Nordheim A. 1999. Id genes are direct targets of bone morphogenetic protein induction in embryonic stem cells. J Biol Chem 274: 19838–19845.

36. Hudson BP, Martinez-Yamout MA, Dyson HJ, Wright PE. 2004. Recognition of the mRNA AU-rich element by the zinc finger domain of TIS11d. Nat Struct Mol Biol 11: 257–264.

37. Imarazene B, Du K, Beille S, Jouanno E, Feron R, Pan Q, Torres-Paz J, Lopez-Roques C, Castinel A, Gil L et al. 2021. A supernumerary “B-sex” chromosome drives male sex determination in the Pachon cavefish, Astyanax mexicanus. Current biology : CB 31: 4800–4809 e4809.

38. Indjeian VB, Kingman GA, Jones FC, Guenther CA, Grimwood J, Schmutz J, Myers RM, Kingsley DM. 2016. Evolving New Skeletal Traits by cis-Regulatory Changes in Bone Morphogenetic Proteins. Cell 164: 45–56.

39. Jiang L, Zhang J, Wang JJ, Wang L, Zhang L, Li G, Yang X, Ma X, Sun X, Cai J et al. 2013. Sperm, but not oocyte, DNA methylome is inherited by zebrafish early embryos. Cell 153: 773–784.

40. Jubb R. 1971. A new Nothobranchius (Pisces, Cyprinodontidae) from Southeastern Rhodesia. Journal of the American Killifish Association 8: 12–19.

41. Kamiya T, Kai W, Tasumi S, Oka A, Matsunaga T, Mizuno N, Fujita M, Suetake H, Suzuki S, Hosoya S et al. 2012. A trans-species missense SNP in Amhr2 is associated with sex determination in the tiger pufferfish, Takifugu rubripes (fugu). PLoS genetics 8: e1002798.

42. Kelmer Sacramento E, Kirkpatrick JM, Mazzetto M, Baumgart M, Bartolome A, Di Sanzo S, Caterino C, Sanguanini M, Papaevgeniou N, Lefaki M et al. 2020. Reduced proteasome activity in the aging brain results in ribosome stoichiometry loss and aggregation. Mol Syst Biol 16: e9596.

43. Kent TV, Uzunovic J, Wright SI. 2017. Coevolution between transposable elements and recombination. Philosophical transactions of the Royal Society of London Series B, Biological sciences 372: 20160458.

44. Kikuchi K, Hamaguchi S. 2013. Novel sex-determining genes in fish and sex chromosome evolution. Developmental dynamics : an official publication of the American Association of Anatomists 242: 339–353.

45. Kikuchi M, Nishimura T, Ishishita S, Matsuda Y, Tanaka M. 2020. foxl3, a sexual switch in germ cells, initiates two independent molecular pathways for commitment to oogenesis in medaka. Proc Natl Acad Sci U S A 117: 12174–12181.

46. Kikuchi M, Nishimura T, Saito D, Shigenobu S, Takada R, Gutierrez-Triana JA, Cerdan JLM, Takada S, Wittbrodt J, Suyama M et al. 2019. Novel components of germline sex determination acting downstream of foxl3 in medaka. Developmental biology 445: 80–89.

47. Kim Y, Nam HG, Valenzano DR. 2016. The short-lived African turquoise killifish: an emerging experimental model for ageing. Dis Model Mech 9: 115–129.

48. Kluver N, Herpin A, Braasch I, Driessle J, Schartl M. 2009. Regulatory back-up circuit of medaka Wt1 co-orthologs ensures PGC maintenance. Developmental biology 325: 179–188.

49. Kluver N, Pfennig F, Pala I, Storch K, Schlieder M, Froschauer A, Gutzeit HO, Schartl M. 2007. Differential expression of anti-Mullerian hormone (amh) and anti-Mullerian hormone receptor type II (amhrII) in the teleost medaka. Developmental dynamics : an official publication of the American Association of Anatomists 236: 271–281.

50. Kobayashi T, Matsuda M, Kajiura-Kobayashi H, Suzuki A, Saito N, Nakamoto M, Shibata N, Nagahama Y. 2004. Two DM domain genes, DMY and DMRT1, involved in testicular differentiation and development in the medaka, Oryzias latipes. Developmental dynamics : an official publication of the American Association of Anatomists 231: 518-526.

51. Krispin S, Stratman AN, Melick CH, Stan RV, Malinverno M, Gleklen J, Castranova D, Dejana E, Weinstein BM. 2018. Growth Differentiation Factor 6 Promotes Vascular Stability by Restraining Vascular Endothelial Growth Factor Signaling. Arteriosclerosis, thrombosis, and vascular biology 38: 353–362.

52. Krug J, Perner B, Albertz C, Morl H, Hopfenmuller VL, Englert C. 2023. Generation of a transparent killifish line through multiplex CRISPR/Cas9mediated gene inactivation. eLife 12: e81549.

53. Kwan KM, Fujimoto E, Grabher C, Mangum BD, Hardy ME, Campbell DS, Parant JM, Yost HJ, Kanki JP, Chien CB. 2007. The Tol2kit: a multisite gateway-based construction kit for Tol2 transposon transgenesis constructs. Developmental dynamics : an official publication of the American Association of Anatomists 236: 3088–3099.

54. Labun K, Montague TG, Krause M, Torres Cleuren YN, Tjeldnes H, Valen E. 2019. CHOPCHOP v3: expanding the CRISPR web toolbox beyond genome editing. Nucleic Acids Res 47: W171–W174.

55. Li L, Dong J, Yan L, Yong J, Liu X, Hu Y, Fan X, Wu X, Guo H, Wang X et al. 2017. Single-Cell RNA-Seq Analysis Maps Development of Human Germline Cells and Gonadal Niche Interactions. Cell Stem Cell 20: 858–873 e854.

56. Li M, Sun Y, Zhao J, Shi H, Zeng S, Ye K, Jiang D, Zhou L, Sun L, Tao W et al. 2015. A Tandem Duplicate of Anti-Mullerian Hormone with a Missense SNP on the Y Chromosome Is Essential for Male Sex Determination in Nile Tilapia, Oreochromis niloticus. PLoS genetics 11: e1005678.

57. Liao Y, Smyth GK, Shi W. 2014. featureCounts: an efficient general purpose program for assigning sequence reads to genomic features. Bioinformatics 30: 923–930.

58. Liao Y, Wang J, Jaehnig EJ, Shi Z, Zhang B. 2019. WebGestalt 2019: gene set analysis toolkit with revamped UIs and APIs. Nucleic Acids Res 47: W199–W205.

59. Liu Y, Kossack ME, McFaul ME, Christensen LN, Siebert S, Wyatt SR, Kamei CN, Horst S, Arroyo N, Drummond IA et al. 2022. Single-cell transcriptome reveals insights into the development and function of the zebrafish ovary. eLife 11: e76014.

60. Love MI, Huber W, Anders S. 2014. Moderated estimation of fold change and dispersion for RNA-seq data with DESeq2. Genome biology 15: 550.

61. Mather JP. 1980. Establishment and characterization of two distinct mouse testicular epithelial cell lines. Biol Reprod 23: 243–252.

62. Mortlock DP, Fang ZM, Chandler KJ, Hou Y, Bickford LR, de Bock CE, Eapen V, Clarke RA. 2022. Transcriptional Interference Regulates the Evolutionary Development of Speech. Genes 13: 1195.

63. Mortlock DP, Guenther C, Kingsley DM. 2003. A general approach for identifying distant regulatory elements applied to the Gdf6 gene. Genome research 13: 2069–2081.

64. Myosho T, Otake H, Masuyama H, Matsuda M, Kuroki Y, Fujiyama A, Naruse K, Hamaguchi S, Sakaizumi M. 2012. Tracing the emergence of a novel sex-determining gene in medaka, Oryzias luzonensis. Genetics 191: 163–170.

64 Nishimura T, Sato T, Yamamoto Y, Watakabe I, Ohkawa Y, Suyama M, Kobayashi S, Tanaka M. 2015. Sex determination. foxl3 is a germ cell-intrinsic factor involved in sperm-egg fate decision in medaka. Science (New York, NY) 349: 328–331.

66. Nishimura T, Tanaka M. 2016. The Mechanism of Germline Sex Determination in Vertebrates. Biol Reprod 95: 30.

67. Nojima H, Rothhamel S, Shimizu T, Kim CH, Yonemura S, Marlow FL, Hibi M. 2010. Syntabulin, a motor protein linker, controls dorsal determination. Development (Cambridge, England) 137: 923–933.

68. Oh D, Houston DW. 2017. Role of maternal Xenopus syntabulin in germ plasm aggregation and primordial germ cell specification. Developmental biology 432: 237–247.

69. Ortega-Recalde O, Hore TA. 2019. DNA methylation in the vertebrate germline: balancing memory and erasure. Essays in biochemistry 63: 649–661.

70. Orvis GD, Jamin SP, Kwan KM, Mishina Y, Kaartinen VM, Huang S, Roberts AB, Umans L, Huylebroeck D, Zwijsen A et al. 2008. Functional redundancy of TGF-beta family type I receptors and receptor-Smads in mediating anti-Mullerian hormone-induced Mullerian duct regression in the mouse. Biol Reprod 78: 994–1001.

71. Pan Q, Feron R, Yano A, Guyomard R, Jouanno E, Vigouroux E, Wen M, Busnel JM, Bobe J, Concordet JP et al. 2019. Identification of the master sex determining gene in Northern pike (Esox lucius) reveals restricted sex chromosome differentiation. PLoS genetics 15: e1008013.

72. Pan Q, Kay T, Depince A, Adolfi M, Schartl M, Guiguen Y, Herpin A. 2021. Evolution of master sex determiners: TGF-beta signalling pathways at regulatory crossroads. Philosophical transactions of the Royal Society of London Series B, Biological sciences 376: 20200091.

73. Peichel CL, McCann SR, Ross JA, Naftaly AFS, Urton JR, Cech JN, Grimwood J, Schmutz J, Myers RM, Kingsley DM et al. 2020. Assembly of the threespine stickleback Y chromosome reveals convergent signatures of sex chromosome evolution. Genome biology 21: 177.

74. Platzer M, Englert C. 2016. Nothobranchius furzeri: A Model for Aging Research and More. Trends in genetics : TIG 32: 543–552.

75. Portnoy ME, McDermott KJ, Antonellis A, Margulies EH, Prasad AB, Program NCS, Kingsley DM, Green ED, Mortlock DP. 2005. Detection of potential GDF6 regulatory elements by multispecies sequence comparisons and identification of a skeletal joint enhancer. Genomics 86: 295–305.

76. Pregizer S, Mortlock DP. 2009. Control of BMP gene expression by long-range regulatory elements. Cytokine & growth factor reviews 20: 509–515.

77. Rácz GA, Nagy N, Tóvári J, Apáti Á, Vértessy BG. 2021. Identification of new reference genes with stable expression patterns for gene expression studies using human cancer and normal cell lines. Scientific Reports 11: 19459.

78. Reed NP, Mortlock DP. 2010. Identification of a distant cis-regulatory element controlling pharyngeal arch-specific expression of zebrafish gdf6a/radar. Developmental dynamics : an official publication of the American Association of Anatomists 239: 1047–1060.

79. Reichwald K, Petzold A, Koch P, Downie BR, Hartmann N, Pietsch S, Baumgart M, Chalopin D, Felder M, Bens M et al. 2015. Insights into Sex Chromosome Evolution and Aging from the Genome of a Short-Lived Fish. Cell 163: 1527–1538.

80. Reuter H, Krug J, Singer P, Englert C. 2018. The African turquoise killifish Nothobranchius furzeri as a model for aging research. Drug Discovery Today: Disease Models 27: 15–22.

81. Richter A, Krug J, Englert C. 2022. Molecular Sexing of Nothobranchius furzeri Embryos and Larvae. Cold Spring Harb Protoc 2022: 630–640.

82. Settle, SH Jr., Rountree RB, Sinha A, Thacker A, Higgins K, Kingsley DM. 2003. Multiple joint and skeletal patterning defects caused by single and double mutations in the mouse Gdf6 and Gdf5 genes. Developmental biology 254: 116–130.

83. Sha QQ, Yu JL, Guo JX, Dai XX, Jiang JC, Zhang YL, Yu C, Ji SY, Jiang Y, Zhang SY et al. 2018. CNOT6L couples the selective degradation of maternal transcripts to meiotic cell cycle progression in mouse oocyte. EMBO J 37.

84. Sharifi-Zarchi A, Gerovska D, Adachi K, Totonchi M, Pezeshk H, Taft RJ, Scholer HR, Chitsaz H, Sadeghi M, Baharvand H et al. 2017. DNA methylation regulates discrimination of enhancers from promoters through a H3K4me1-H3K4me3 seesaw mechanism. BMC Genomics 18: 964.

85. Smith T, Heger A, Sudbery I. 2017. UMI-tools: modeling sequencing errors in Unique Molecular Identifiers to improve quantification accuracy. Genome research 27: 491–499.

86. Stundlova J, Hospodarska M, Luksikova K, Volenikova A, Pavlica T, Altmanova M, Richter A, Reichard M, Dalikova M, Pelikanova S et al. 2022. Sex chromosome differentiation via changes in the Y chromosome repeat landscape in African annual killifishes Nothobranchius furzeri and N. kadleci. Chromosome Res 30: 309–333.

87. Tanaka M. 2016. Germline stem cells are critical for sexual fate decision of germ cells. Bioessays 38: 1227–1233.

88. Tassabehji M, Fang ZM, Hilton EN, McGaughran J, Zhao Z, de Bock CE, Howard E, Malass M, Donnai D, Diwan A et al. 2008. Mutations in GDF6 are associated with vertebral segmentation defects in Klippel-Feil syndrome. Human mutation 29: 1017–1027.

89. Terzibasi E, Valenzano DR, Benedetti M, Roncaglia P, Cattaneo A, Domenici L, Cellerino A. 2008. Large differences in aging phenotype between strains of the short-lived annual fish Nothobranchius furzeri. PloS one 3: e3866.

90. Thoma EC, Wagner TU, Weber IP, Herpin A, Fischer A, Schartl M. 2011. Ectopic expression of single transcription factors directs differentiation of a medaka spermatogonial cell line. Stem Cells Dev 20: 1425–1438.

91. Tozzini ET, Dorn A, Ng’oma E, Polacik M, Blazek R, Reichwald K, Petzold A, Watters B, Reichard M, Cellerino A. 2013. Parallel evolution of senescence in annual fishes in response to extrinsic mortality. BMC Evol Biol 13: 77.

92. Valenzano DR, Benayoun BA, Singh PP, Zhang E, Etter PD, Hu CK, Clément-Ziza M, Willemsen D, Cui R, Harel I et al. 2015. The African Turquoise Killifish Genome Provides Insights into Evolution and Genetic Architecture of Lifespan. Cell 163: 1539–1554.

93. Valenzano DR, Kirschner J, Kamber RA, Zhang E, Weber D, Cellerino A, Englert C, Platzer M, Reichwald K, Brunet A. 2009. Mapping loci associated with tail color and sex determination in the short-lived fish Nothobranchius furzeri. Genetics 183: 1385–1395.

94. Venkatesan AM, Vyas R, Gramann AK, Dresser K, Gujja S, Bhatnagar S, Chhangawala S, Gomes CBF, Xi HS, Lian CG et al. 2018. Ligand-activated BMP signaling inhibits cell differentiation and death to promote melanoma. The Journal of clinical investigation 128: 294–308.

95. Wang J, Yu T, Wang Z, Ohte S, Yao RE, Zheng Z, Geng J, Cai H, Ge Y, Li Y et al. 2016. A New Subtype of Multiple Synostoses Syndrome Is Caused by a Mutation in GDF6 That Decreases Its Sensitivity to Noggin and Enhances Its Potency as a BMP Signal. Journal of bone and mineral research : the official journal of the American Society for Bone and Mineral Research 31: 882–889.

96. Wang R, Liu X, Li L, Yang M, Yong J, Zhai F, Wen L, Yan L, Qiao J, Tang F. 2022. Dissecting Human Gonadal Cell Lineage Specification and Sex Determination Using A Single-cell RNA-seq Approach. Genomics Proteomics Bioinformatics 20: 223–245.

97. Wu Q, Fukuda K, Weinstein M, Graff JM, Saga Y. 2015. SMAD2 and p38 signaling pathways act in concert to determine XY primordial germ cell fate in mice. Development (Cambridge, England) 142: 575–586.

98. Yang J, Li X, Li Y, Southwood M, Ye L, Long L, Al-Lamki RS, Morrell NW. 2013. Id proteins are critical downstream effectors of BMP signaling in human pulmonary arterial smooth muscle cells. American journal of physiology Lung cellular and molecular physiology 305: L312–321.

99. Yano A, Guyomard R, Nicol B, Jouanno E, Quillet E, Klopp C, Cabau C, Bouchez O, Fostier A, Guiguen Y. 2012. An immune-related gene evolved into the master sex-determining gene in rainbow trout, Oncorhynchus mykiss. Current biology : CB 22: 1423–1428.

100. Zhu A, Ibrahim JG, Love MI. 2019. Heavy-tailed prior distributions for sequence count data: removing the noise and preserving large differences. Bioinformatics 35: 2084–2092.

